# Structure of *Escherichia coli* respiratory complex I reconstituted into lipid nanodiscs reveals an uncoupled conformation

**DOI:** 10.1101/2021.04.09.439197

**Authors:** Piotr Kolata, Rouslan G. Efremov

**Affiliations:** Center for Structural Biology, Vlaams Instituut voor Biotechnologie, Brussels, Belgium; Structural Biology Brussels, Department of Bioengineering Sciences, Vrije Universiteit Brussel, Brussels, Belgium

## Abstract

Respiratory complex I is a multi-subunit membrane protein complex that reversibly couples NADH oxidation and ubiquinone reduction with proton translocation against trans-membrane potential. Complex I from *Escherichia coli* is among the best functionally characterized complexes, but its structure remains unknown, hindering further mechanistic studies to understand the enzyme coupling mechanism. Here we describe the single particle cryo-electron microscopy (cryo-EM) structure of the entire catalytically active *E. coli* complex I reconstituted into lipid nanodiscs. The structure of this mesophilic bacterial complex I displays highly dynamic connection between the peripheral and membrane domains. The peripheral domain assembly is stabilized by unique terminal extensions and an insertion loop. The membrane domain structure reveals novel dynamic features. Unusual conformation of the conserved interface between the cytoplasmic and membrane domains suggests an uncoupled conformation of the complex. Based on these structural data we suggest a new simple and testable coupling mechanism for the molecular machine.

## Introduction

Complex I, NADH:ubiquinone oxidoreductase, is a multi-subunit enzyme found in many bacteria and most eukaryotes. It facilitates transfer of two electrons from NADH to ubiquinone, or its analogues, coupled reversibly with translocation of four protons across the membrane against trans-membrane potential (Galkin et al., 2006; Sazanov, 2015). Structures of the complete complex I from several eukaryotes (Fiedorczuk et al., 2016; Hunte et al., 2010; Kampjut and Sazanov, 2020; Zhu et al., 2016), one thermophilic bacterium (Baradaran et al., 2013), and the partial structure of the membrane domain of *Escherichia coli* complex I (Efremov and Sazanov, 2011), have been determined.

The composition of complex I differs significantly between species. Mitochondrial complex I has molecular weight 1 MDa and comprises more than 35 subunits (Wirth et al., 2016) whereas bacterial analogues are much smaller with molecular weights approximately 500 kDa. Complex I from all characterized species contains homologues of 14 core subunits; seven subunits each assemble into peripheral and membrane arms, joined at their tips and form the complex with a characteristic L-shape.

The peripheral arm, exposed to the cytoplasm in bacteria or the mitochondrial matrix in eukaryotes, contains binding sites for NADH, ubiquinone, and flavin mononucleotide (FMN) as well as eight or nine iron-sulfur clusters, seven of which connect the NADH and ubiquinone-binding sites (Sazanov, 2015) enabling rapid electron transfer (Verkhovskaya et al., 2008) .

The membrane-embedded arm includes a chain of three antiporter-like subunits, NuoL, NuoM, and NuoN (*E. coli* nomenclature is used for the subunits hereafter) (Efremov and Sazanov, 2011), which are also found in the Mrp family of multisubunit H^+^/Na antiporters (Steiner and Sazanov, 2020). Each antiporter-like subunit contains two structural repeats comprising five trans-membrane helices (TMH, TMH4-8, and TMH9-13). TMH7 and TMH12 are interrupted by an extended loop in the middle of the membrane and the helix TM8 at the interface between symmetric motifs is interrupted by the *π*-bulge (Baradaran et al., 2013; Efremov and Sazanov, 2011). Membrane-embedded NuoH mediates interaction with the peripheral arm and also contains five-helix structural repeats found in antiporter-like subunits (Baradaran et al., 2013). Together with subunits NuoB and NuoD it forms an extended ubiquinone-binding cavity (Q-cavity) spanning the membrane bilayer hydrophobic region to the ubiquinone-binding site (Q-site) in the proximity of the terminal iron-sulfur cluster N2 (Baradaran et al., 2013).

The membrane arm features a continuous chain of conserved and functionally important ionizable residues positioned in the middle of the membrane. These are suggested to be involved in proton translocation and its coupling to electron transfer (Baradaran et al., 2013; Efremov and Sazanov, 2011). Attempts to visualize conformational changes in the membrane domain (Kampjut and Sazanov, 2020; Parey et al., 2018) have revealed rotation of the cytoplasmic half of TMH3 of NuoJ in mammalian complex I (Agip et al., 2018) and were associated with active-deactive transition. Recently, proton translocation mechanisms without conformational changes in antiporter-like subunits were suggested (Kampjut and Sazanov, 2020; Steiner and Sazanov, 2020). However, all proposed coupling mechanisms remain largely speculative and require further validation by functional, biochemical, and structural methods.

*E. coli* complex I is among the best functionally characterized complex I. It has been studied using many biophysical and biochemical techniques (Verkhovskaya and Bloch, 2012). Combined with the possibility of fast and extensive mutagenesis (Pohl et al., 2007; Verkhovskaya and Bloch, 2012), it represents a highly attractive system to study the coupling mechanism. However, owing to its fragile and dynamic nature (Verkhovskaya and Bloch, 2012), high-resolution structures of this complex remain limited to a partial structure of the membrane domain (Efremov and Sazanov, 2011).

Here we present a single particle cryo-EM structure of the entire *E. coli* complex I reconstituted into lipid nanodiscs, with the peripheral arm structure solved at 2.1 Å resolution and that of the membrane domain at 3.7 Å.

## Results

### Overall structure

Twin-strep tag was added to genomically encoded subunit NuoF using a CRISPR-Cas9 based system (Jiang et al., 2015) (Figure 1 - figure supplement 1). This enabled single-step purification of solubilized complex (Figure 1 - figure supplement 2A), which was further reconstituted into lipid nanodiscs comprising *E. coli* polar lipids and membrane scaffold protein MSP2N2 (Grinkova et al., 2010) (Figure 1 - figure supplement 2A,B). Mass photometry indicated that reconstituted complex I was homogeneous and monodispersed (Figure 1 - figure supplement 2C,D).

**Figure 1.**
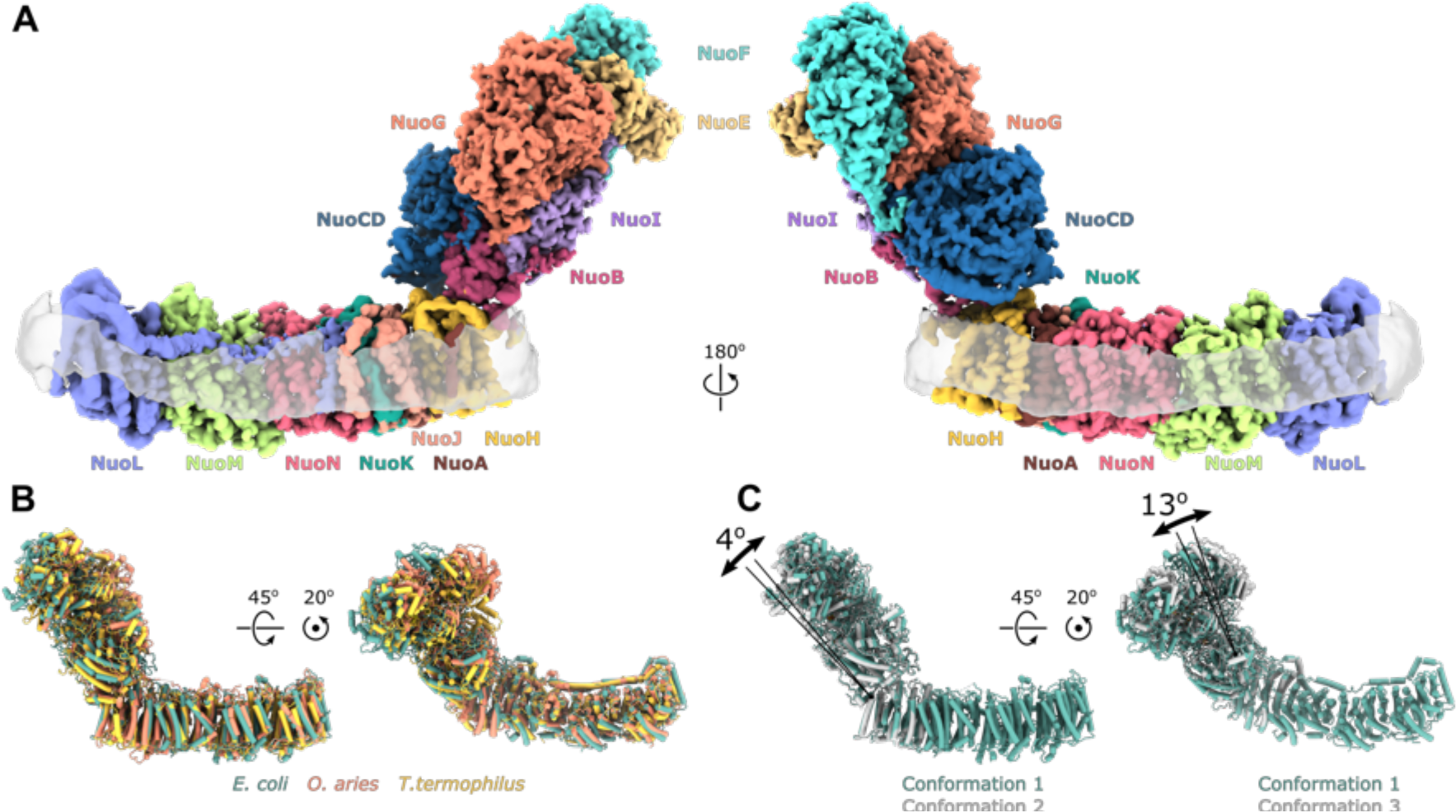
Architecture of *Escherichia coli* respiratory complex I. (A) Segmented density map of the complete complex I shown together with the nanodisc density. (B) Comparison of the structures of the *E. coli* (green), *Thermus thermophilus* (PDB ID: 4HEA, yellow), and the core subunits of ovine (PDB ID: 6ZKD, orange) complex I. (C) Conformational differences between three conformations resolved at high resolution. The structures are aligned on the membrane arm. The rotation axes and angles are indicated.

NADH:potassium ferricyanide (FeCy) and NADH:ubiquinone-1 (Q1) activities of the reconstituted complex I (Figure 1 - figure supplement 2E,F) were similar to those of detergent-purified protein supplemented with native *E. coli* lipids (Sazanov, 2003).

Furthermore, NADH:Q1 activity was completely inhibited by piericidin-A (Figure 1 - figure supplement 2F) indicating that complex I reconstituted in lipid nanodiscs was intact and catalytically active in a detergent-free environment.

We determined the single particle cryo-EM structure of the reconstituted complex (Figure 1, Figure 1 - figure supplement 3,4, Table 1, Movie 1). Multiple conformations of the complex that differed by relative positions of the peripheral and membrane arms were revealed by 3D classification (Figure 1 - figure supplement 4,5). Three conformations of the entire complex were reconstructed to average resolutions between 3.3 and 3.7 Å (Figure 1 - figure supplement 4) resolving the interface between the arms; however, due to high-residual mobility of the arms, the antiporter-like subunits were resolved at below 8 Å (Figure 1 - figure supplement 4).

**Table 1.**
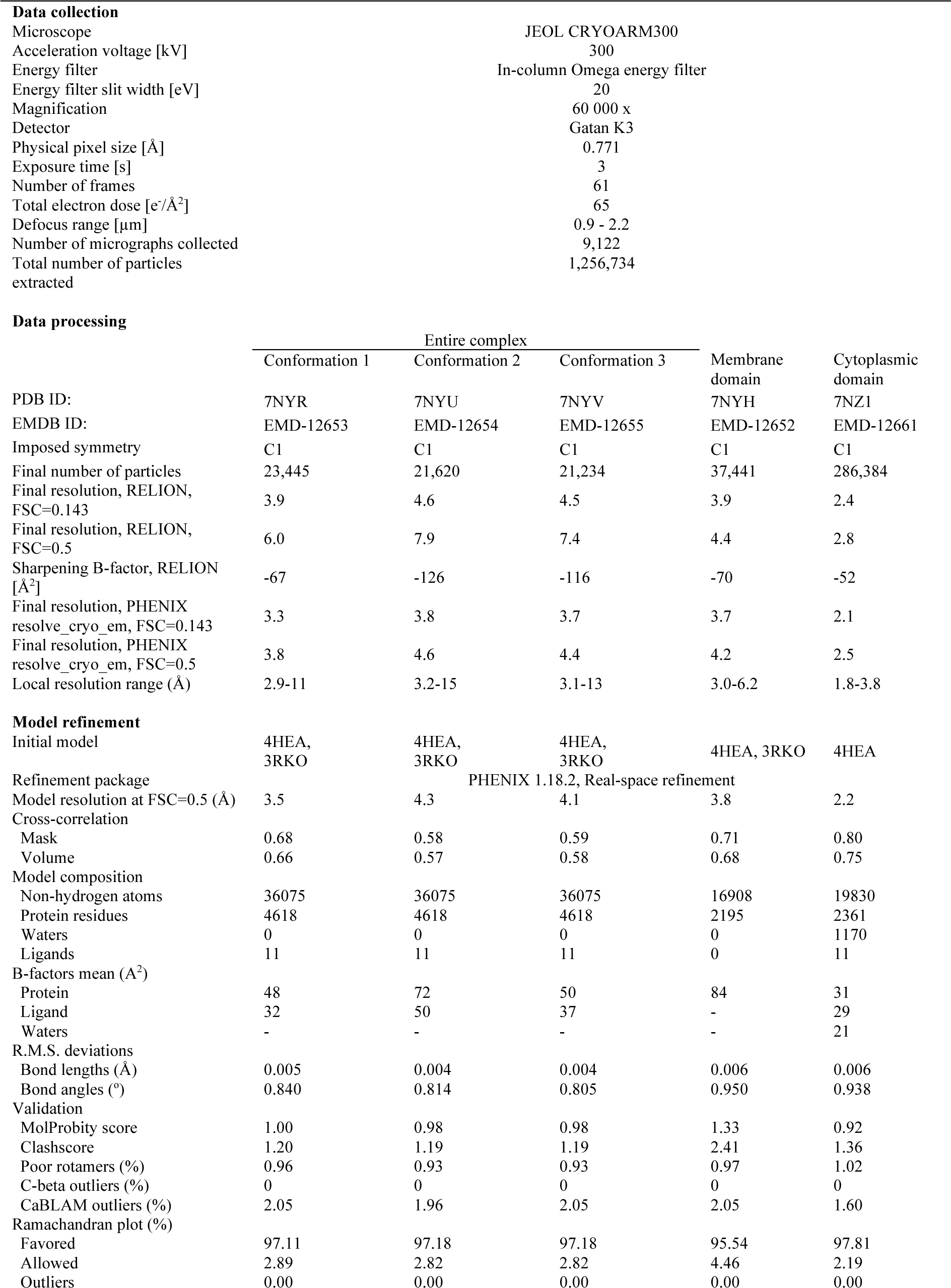
Statistics of cryo-EM data collection, data processing, and model refinement

| Data collection |  |
| --- | --- |
| Microscope | JEOL CRYOARM300 |
| Acceleration voltage [kV] | 300 |
| Energy filter | In-column Omega energy filter |
| Energy filter slit width [eV] | 20 |
| Magnification | 60 000 x |
| Detector | Gatan K3 |
| Physical pixel size [Å] | 0.771 |
| Exposure time [s] | 3 |
| Number of frames | 61 |
| Total electron dose [e<sup>-</sup>/Å<sup>2</sup>] | 65 |
| Defocus range [μm] | 0.9 - 2.2 |
| Number of micrographs collected | 9,122 |
| Total number of particles extracted | 1,256,734 |

| Data processing |  |  |  |  |  |
| --- | --- | --- | --- | --- | --- |
|  | Entire complex |  |  |  |  |
|  | Conformation 1 | Conformation 2 | Conformation 3 | Membrane domain | Cytoplasmic domain |
| PDB ID: | 7NYR | 7NYU | 7NYV | 7NYH | 7NZ1 |
| EMDB ID: | EMD-12653 | EMD-12654 | EMD-12655 | EMD-12652 | EMD-12661 |
| Imposed symmetry | C1 | C1 | C1 | C1 | C1 |
| Final number of particles | 23,445 | 21,620 | 21,234 | 37,441 | 286,384 |
| Final resolution, RELION, FSC=0.143 | 3.9 | 4.6 | 4.5 | 3.9 | 2.4 |
| Final resolution, RELION, FSC=0.5 | 6.0 | 7.9 | 7.4 | 4.4 | 2.8 |
| Sharpening B-factor, RELION [Å<sup>2</sup>] | -67 | -126 | -116 | -70 | -52 |
| Final resolution, PHENIX resolve_cryo_em, FSC=0.143 | 3.3 | 3.8 | 3.7 | 3.7 | 2.1 |
| Final resolution, PHENIX resolve_cryo_em, FSC=0.5 | 3.8 | 4.6 | 4.4 | 4.2 | 2.5 |
| Local resolution range (Å) | 2.9-11 | 3.2-15 | 3.1-13 | 3.0-6.2 | 1.8-3.8 |

| Model refinement |  |  |  |  |  |
| --- | --- | --- | --- | --- | --- |
| Initial model | 4HEA, 3RKO | 4HEA, 3RKO | 4HEA, 3RKO | 4HEA, 3RKO | 4HEA |
| Refinement package | PHENIX 1.18.2, Real-space refinement |  |  |  |  |
| Model resolution at FSC=0.5 (Å) | 3.5 | 4.3 | 4.1 | 3.8 | 2.2 |
| Cross-correlation |  |  |  |  |  |
| Mask | 0.68 | 0.58 | 0.59 | 0.71 | 0.80 |
| Volume | 0.66 | 0.57 | 0.58 | 0.68 | 0.75 |
| Model composition |  |  |  |  |  |
| Non-hydrogen atoms | 36075 | 36075 | 36075 | 16908 | 19830 |
| Protein residues | 4618 | 4618 | 4618 | 2195 | 2361 |
| Waters | 0 | 0 | 0 | 0 | 1170 |
| Ligands | 11 | 11 | 11 | 0 | 11 |
| B-factors mean (Å<sup>2</sup>) |  |  |  |  |  |
| Protein | 48 | 72 | 50 | 84 | 31 |
| Ligand | 32 | 50 | 37 | - | 29 |
| Waters | - | - | - | - | 21 |
| R.M.S. deviations |  |  |  |  |  |
| Bond lengths (Å) | 0.005 | 0.004 | 0.004 | 0.006 | 0.006 |
| Bond angles (°) | 0.840 | 0.814 | 0.805 | 0.950 | 0.938 |
| Validation |  |  |  |  |  |
| MolProbity score | 1.00 | 0.98 | 0.98 | 1.33 | 0.92 |
| Clashscore | 1.20 | 1.19 | 1.19 | 2.41 | 1.36 |
| Poor rotamers (%) | 0.96 | 0.93 | 0.93 | 0.97 | 1.02 |
| C-beta outliers (%) | 0 | 0 | 0 | 0 | 0 |
| CaBLAM outliers (%) | 2.05 | 1.96 | 2.05 | 2.05 | 1.60 |
| Ramachandran plot (%) |  |  |  |  |  |
| Favored | 97.11 | 97.18 | 97.18 | 95.54 | 97.81 |
| Allowed | 2.89 | 2.82 | 2.82 | 4.46 | 2.19 |
| Outliers | 0.00 | 0.00 | 0.00 | 0.00 | 0.00 |

Focused refinement of each arm separately and subtraction of nanodisc density (Figure 1 - figure supplement 3) improved the resolution of peripheral and membrane arms to 2.7 Å and 3.7 Å, respectively (Figure 1 - figure supplement 3,4, Table 1).

Micrograph analysis, in contrast to mass photometry, revealed that large fraction of the particles corresponds to the peripheral arm only (Figure 1 - figure supplement 3) that may have dissociated during cryo-EM sample preparation. These yielded 3D reconstruction to 2.8 Å resolution (Figure 1 - figure supplement 3), similar to the map of the peripheral arm of intact complex I. Joining two subsets improved resolution of the peripheral arm to 2.1 Å (Table 1, Figure 1 - figure supplement 4). Using the resulting maps, an atomic model of the entire *E. coli* complex I was built, comprising 4618 residues and accounting for 94.7% of the total polypeptide constituting the complex (Table 2).

**Table 2.** Residues built in the models

| Subunit | Total<br>number of<br>residues | Built residues |  |  | Co-factors | Fragments build in<br>entire complex only |
| --- | --- | --- | --- | --- | --- | --- |
|  |  | Entire<br>complex | Membrane<br>arm | Cytoplasmic<br>arm |  |  |
| NuoF | 445 | 1-441 |  | 1-441 | FMN, N3 |  |
| NuoE | 166 | 11-166 |  | 11-166 | N1a |  |
| NouG | 908 | 1-907 |  | 1-907 | N1b, N4,<br>N7 |  |
| NuoI | 180 | 23-180 |  | 39-180 | N6a, N6b | 23-38 |
| NuoB | 220 | 43-76,<br>86-179,<br>190-220 |  | 53-71,<br>90-179,<br>190-220 | N2 | 43-53, 71-76, 86-90: |
| NuoCD | 596:<br>C 1-172<br>D 212-596 | 9-596 |  | 9-205,<br>210-218,<br>224-233,<br>238-596 | - | 206-209,<br>219-223,<br>234-237 |
| NuoH | 325 | 52-321 | 52-214,<br>223-321 |  | - | 215-222 |
| NuoA | 147 | 15-38,<br>60-127 | 15-38,<br>66-127 |  | - | 61-66 |
| NuoJ | 184 | 1-164 | 1-164 |  | - |  |
| NuoK | 100 | 1-100 | 1-100 |  | - |  |
| NuoN | 485 | 1-191,<br>199-437,<br>447-483 | 1-191,<br>199-437,<br>447-483 |  | - |  |
| NuoM | 509 | 1-504 | 1-504 |  | - |  |
| NuoL | 613 | 1-612 | 1-612 |  | - |  |

Arrangement of the arms and individual core subunits in *E. coli* complex I is very similar to that of *Thermus thermophilus* (RMSD 6.3 Å over 2593 C*α* atoms, 4.0 Å membrane arm over 1604 atoms, and 2.2 Å for peripheral arm 855 C*α*) and mammalian enzyme (8.3 Å over 2420 C*α* atoms (3.8 Å MD 1710 atoms, PD 2.0 767 C*α*)) (Figure 1B) apart from the relative long-range twisting and bending of arms observed between complex I from different species (Baradaran et al., 2013; Vinothkumar et al., 2014).

Comparison of *E. coli* complex I conformations reconstructed to better than 4 Å resolution revealed two modes of relative arm rotation (Figure 1C): 1) rotation around an axis that passes through the NuoH-NuoB interface and is tilted around 45 degrees out of the plane formed by the arms with an amplitude of at least 13 degrees, and 2) rotation around an axis parallel to the membrane and roughly perpendicular to the long axis of the membrane arm with an amplitude of approximately 4 degrees. Although relative arm movements were observed in mammalian (Kampjut and Sazanov, 2020; Zhu et al., 2016) and *T. thermophilus* complex I (Gutiérrez-Fernández et al., 2020), their amplitudes were smaller and movement directionality was less diverse. Despite significant relative arm movements, the structure of each arm was rigid and did not reveal different conformations apart from the specific local dynamics discussed below.

### Structure of the peripheral arm

#### Architecture of the peripheral arm reveals a novel evolutionary strategy to stabilize the subcomplex

At an average resolution of 2.1 Å with the local resolution in the core reaching 2.0 Å (Figure 1 - figure supplement 4) conformations of most side chains in the peripheral arm, positions of ions, and multiple water molecules were resolved unambiguously (Figure 2, Figure 1 - figure supplement 6).

**Figure 2.**
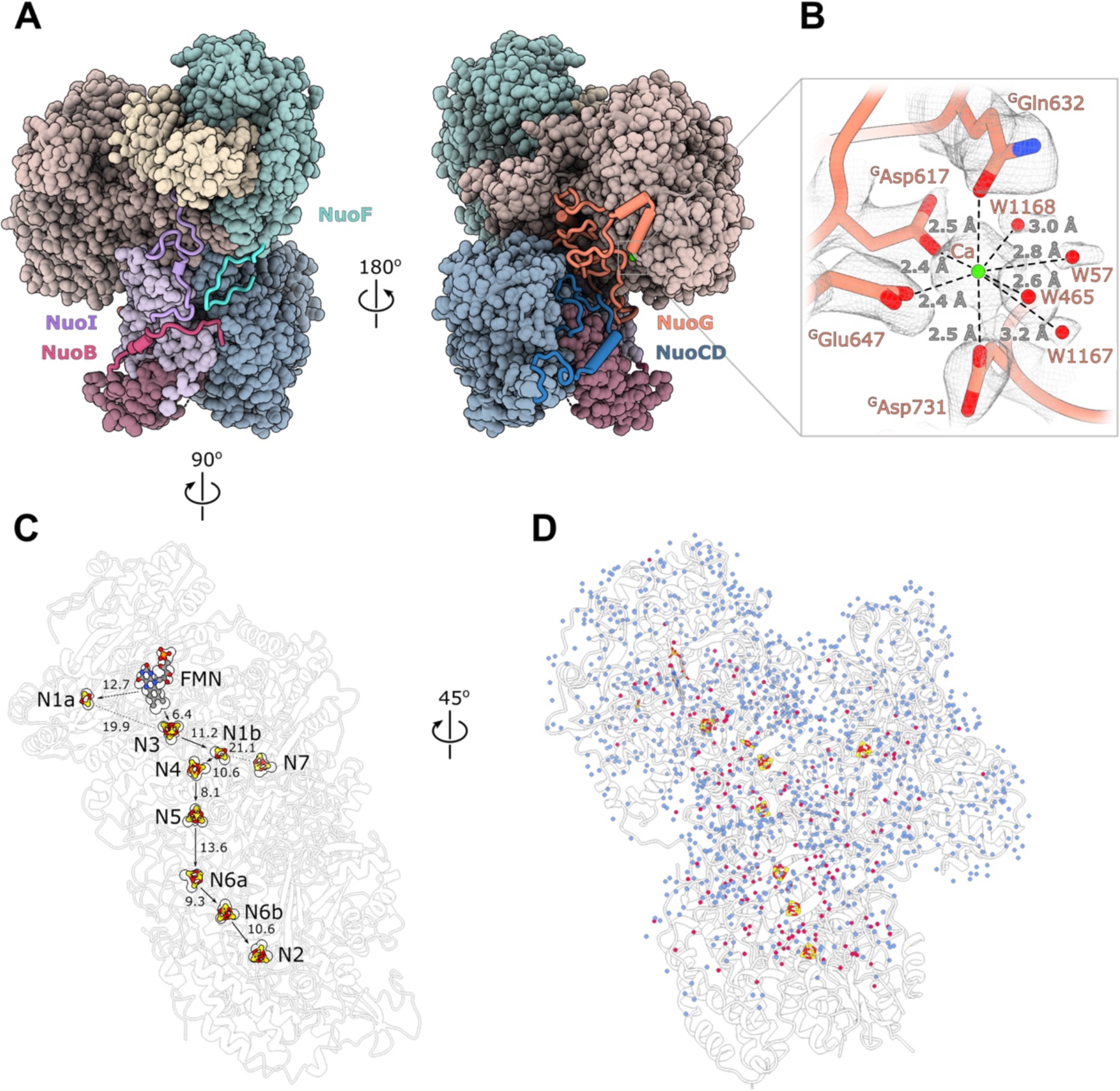
Structure of the peripheral arm. (A, B) The *Escherichia coli-*specific extensions in the peripheral arm subunits: (A) C-termini of NuoI (violet), NuoB (pink), NuoF (turquoise), (B) NuoG insertion (orange), and NuoCD linker (blue). (C) Structural details of the calcium-binding site. (D) Comparison of the FMN and Fe-S clusters positions in *E. coli* (shown as atoms) and *Thermus thermophilus* (shown as outline around *E. coli* atoms). Edge- to-edge distances and the electron pathway are indicated. (E) Water molecules modelled into the 2.1 Å resolution density of the peripheral arm are show in blue. Water molecules conserved with the peripheral arm of ovine complex I (PDB ID: 6ZK9, red) are shown as red spheres. FMN and iron-sulfur (Fe-S) clusters are shown as spheres.

The overall architecture of the conserved core of the peripheral arm subunits is very similar to other homologues. Unlike other structurally characterized homologues, *E. coli* subunits NuoC and NuoD are joined in a single polypeptide. The 35 amino acid-long linker includes an *α*- helix (residues 180-194) that interacts with subunit NuoB (Figure 2A). The relative positions of all redox centers with FMN and nine iron-sulfur clusters, including off path cluster N7 (Sazanov, 2006), are particularly well conserved (Figure 2C).

A distinctive feature of the *E. coli* peripheral arm is the presence of ordered C-terminal extensions in subunits NuoB, NuoI, and NuoF with a length of 22 to 45 residues and a large 94 residue insertion loop in subunit NuoG, referred to as the G-loop (Figure 2A, Table 3). These extensions are unique among structurally characterized complex I homologues and have a well-defined structure. While the G-loop has a compact fold, the conformation of the C-terminal tails is extended. They line the surface of the conserved fold of the peripheral arm with high shape complementarity (Figure 2A, Figure 2 - figure supplement 1). Apart from a few helical turns, these extensions have no secondary structure elements (Table 3). They create additional inter-subunit contacts with some surface areas exceeding 1,000 Å^2^ and involving polar interactions (Table 3). Similarly, the G-loop fills a crevice between NuoCD, NuoI, and NuoG subunits (Figure 2A). Together the extensions and G-loop increase the interaction surface between the electron acceptor module (NuoEFG) and connecting module (NuoICDB) by a factor of three (from 1400 to 4600 Å^2^), thus stabilizing the peripheral arm assembly. These structural features are conserved within the Enterobacteriaceae family and are very common in the phylum Gammaproteobacteria. They display high conservation of interfacial residues, particularly for the G-loop (Figure 2 - figure supplement 1) and demonstrate a new evolutionary strategy for complex stabilization that was not observed to date in complex I structures from other species.

**Table 3.** Properties of *E. coli* peripheral arm extensions (analyzed in PIZA)

| Subunit | Extension residues numbers | Interacts with subunit | Interaction surface, [Å<sup>2</sup>] | Secondary structure | Specific interactions | Spatial overlap with subunits in other species |
| --- | --- | --- | --- | --- | --- | --- |
| NuoF | C-term<br>424-445 | NuoI,<br>NuoD<br>NuoB | 254<br>557<br>118 | no | Nb 8 Sb 0<br>Hb 0 Sb 1<br>Hb 0 Sb 0 | Nqo15 *T. Therophilus*<br>NUIM,NUZM, **NUMM** *,<br>*Y.Lipolytica*<br>**Ndufs6**, Ndufs8, mouse |
| NuoG | Insertion<br>687-781 | NuoCD<br>NuoI | 1080<br>585 | 2 helical<br>turns | Hb 12 Sb 5<br>Hb 13 Sb 4 | Nqo5 *T. Therophilus*<br>NUGM, NUYM<br>*Y.Lipolytica*<br>Ndufs3, Ndufs4 mouse |
| NuoI | C-term<br>139-180 | NuoG<br>NuoB<br>NuoF<br>NuoD<br>NuoE | 805<br>348<br>292<br>122<br>421 | 1 helical<br>turn | Nb 8 Sb 5<br>Nb 1 Sb 0<br>Nb 3 Sb 0<br>Nb 0 Sb 1<br>Nb 1 Sb 0 | Nqo15 *T. Therophilus*<br>**NUMM** *Y.Lipolytica*<br>**Ndufs6** mouse |
| NuoB | C-term<br>196-220 | NuoI<br>NuoD<br>NuoF | 1095<br>216<br>117 | 2 helical<br>turns | Nb 16 Sb 4<br>Nb 1 Sb 3<br>Ng 0 Sb 0 | NUIM, N7BM *Y.Lipolytica*<br>Ndufs8, Ndufa12 mouse |

\*Subunits in bold are equivalent to NuoI

A strong density near the NuoG surface coordinated by ^G^Asp617, ^G^Gln632, ^G^Glu647, ^G^Asp731 and four water molecules (Figure 2B) was assigned to a Ca^2+^ ion. The coordination number, geometry, and ion-ligand distances of ca 2.5 Å (H. Zheng et al., 2008) as well as the 2 mM concentration of Ca^2+^ in the buffer support this assignment. Divalent ions are known to increase both the activity and stability of *E. coli* complex I (Sazanov, 2003). One of the calcium ligands, ^G^Asp731, is part of the G-loop, suggesting that Ca^2+^ stabilizes the fold of the G-loop and consequently, the peripheral arm.

The extensions spatially overlap with the supernumerary subunits of complex I from *T. thermophilus* (Sazanov, 2006) and the structurally conserved supernumerary subunits of eukaryotic complex I (Zhu et al., 2016) (Table 3), consistent with the suggestion that the primary role of supernumerary subunits is to stabilize the complex (Fiedorczuk et al., 2016).

#### Bound water molecules

At 2.1 Å resolution, 1165 water molecules associated with the peripheral arm were modelled (Figure 2D). The positions of 180 water molecules are conserved with those identified in the peripheral arm of ovine complex I (Kampjut and Sazanov, 2020) (Figure 2D, red spheres).

Most of conserved waters are buried in the interior of the subunits, shielded from the solvent, and most likely play a structural role in maintaining the subunit fold. Only a few water molecules interact closely with iron-sulfur clusters and may influence their potential (Table 4, Figure 1 - figure supplement 5). The water molecules located close to or between iron-sulfur clusters are not more conserved than those in the other parts of the complex, suggesting that they were not evolutionary selected to optimize the rate of electron transfer as was suggested by Schulte et.al. (Schulte et al., 2019a).

**Table 4.**
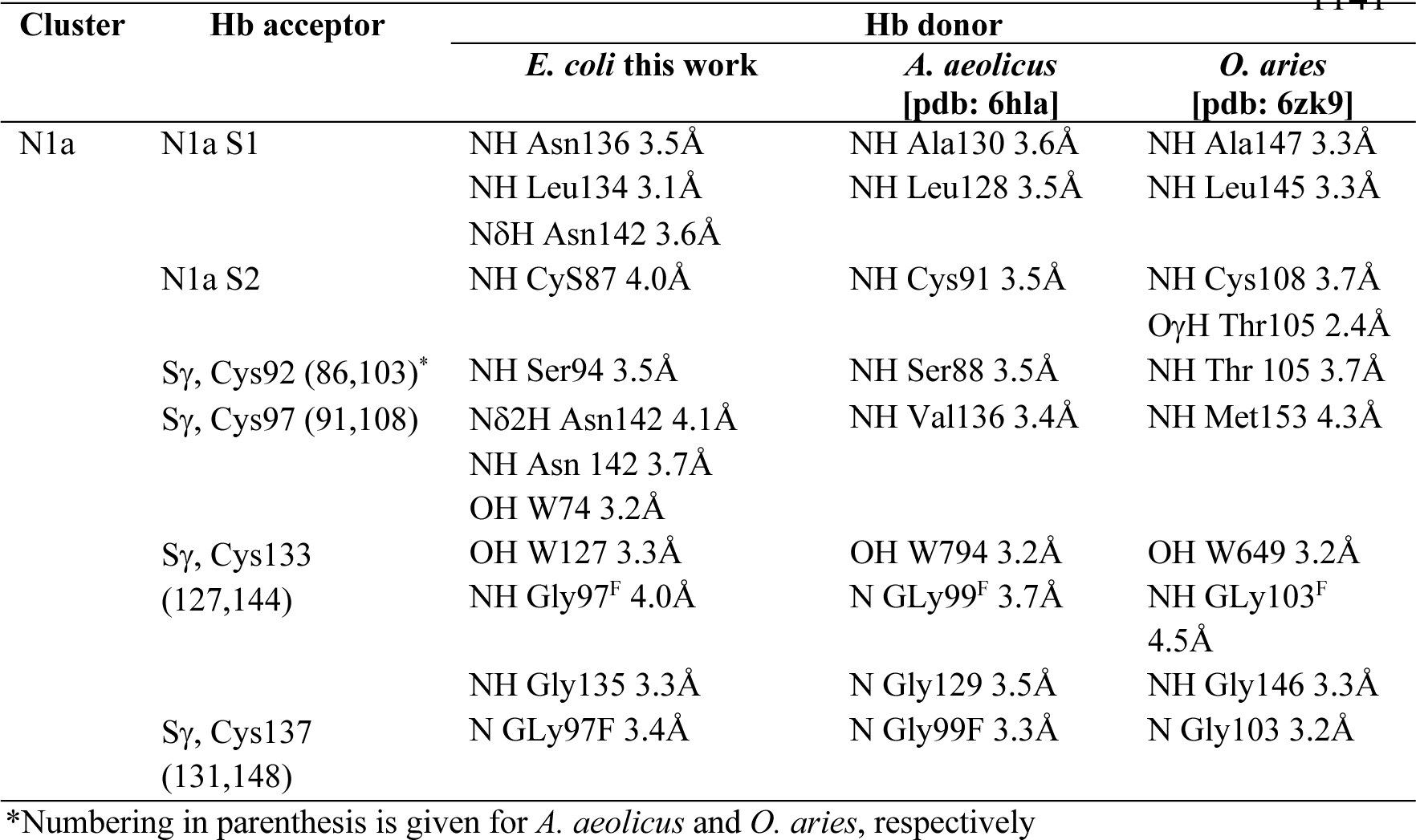
Comparison of hydrogen bond networks surrounding the N1a cluster in complex I structures solved at high resolution

At 2.1 Å resolution, several unusual density features were observed next to some surface-exposed histidines and between some cysteine-methionine pairs as listed in Table 6 and depicted in Figure 3 - figure supplement 1.

**Figure 3.**
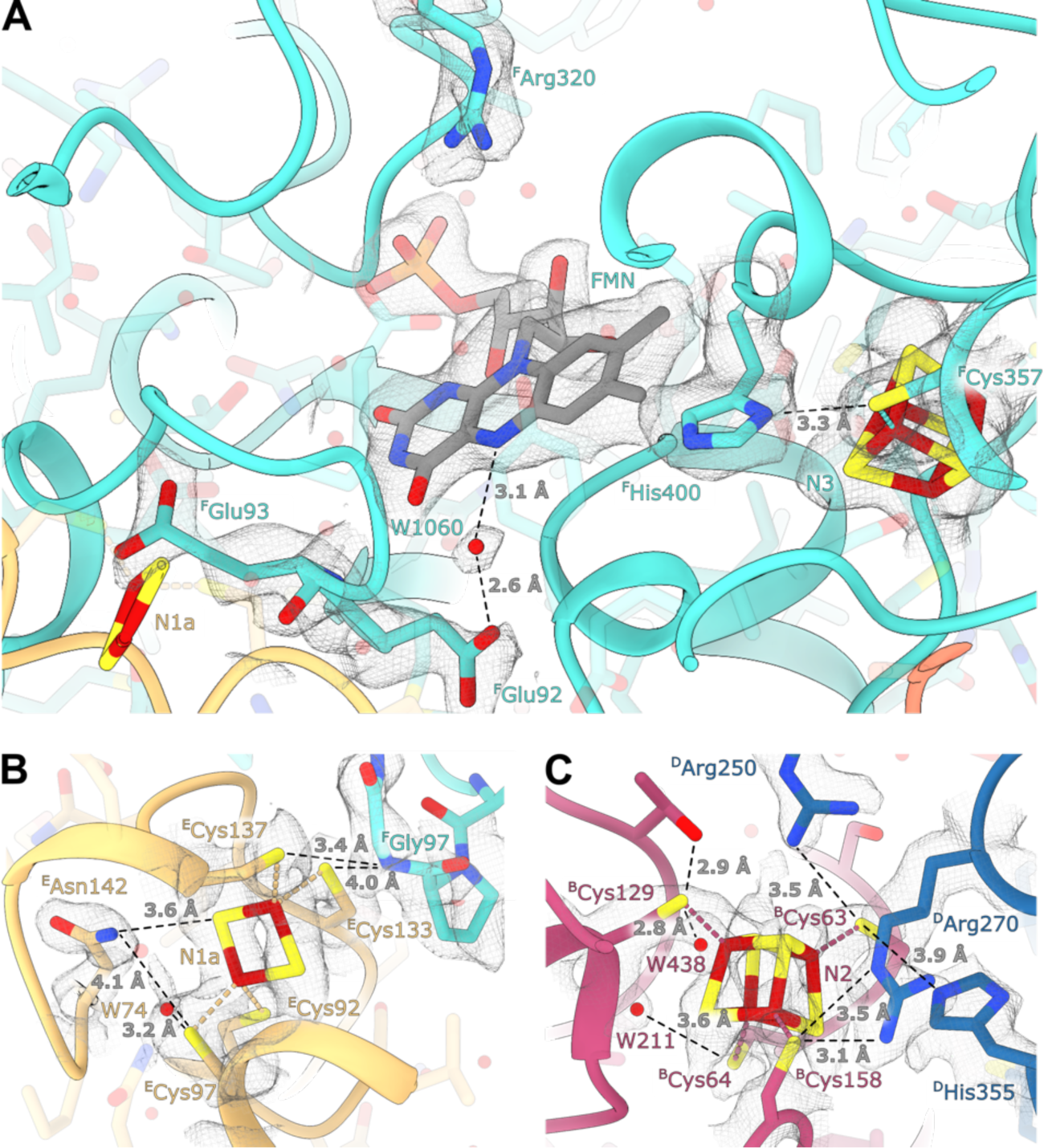
Details of the electron transport chain. (A) The NADH-binding pocket and environment of the Fe-S cluster N3. (B, C) Environment of the Fe-S clusters N1a and N2.

#### Electron input and output sites

The FMN conformation and key water molecules in the NADH-binding pocket of *E. coli* complex I are conserved (Kampjut and Sazanov, 2020; Schulte et al., 2019b). This includes the position of W1060 that forms a hydrogen bond with the isoalloxazine ring N5 atom in FMN and with ^F^Glu92, which likely acts as the activating group during catalysis of hydride transfer from NADH (Fraaije et al., 2000) (Figure 3A). Schulte et.al. (Schulte et al., 2019b) suggested a mechanism for regulation of reactive oxygen species (ROS) generation by *E. coli* complex I that involves flipping the carbonyl oxygen of ^F^Glu93 upon enzymatic reduction.

Our structure unambiguously places the corresponding carbonyl oxygen in a conformation that points away from FMN (Figure 3A) similar to conformations found in the reduced and oxidized ovine complex I (Kampjut and Sazanov, 2020), which does not support its involvement in ROS regulation.

*E. coli*-specific features in the FMN-binding pocket include ^F^His400 that replaces the Leu residues found in other homologues. ^F^His400 is in Van der Waals contact with the isoalloxazine ring of FMN; its imidazole ring interacts directly with the N3 cluster iron atom and forms a hydrogen bond with S*γ* of ^F^Cys357 coordinating N3 (Figure3A). ^F^His400 is solvent-accessible even in the presence of NADH, and therefore, may become protonated upon N3 reduction. ^F^Arg320 is positioned such that it can form hydrogen bonds with the ribose moiety of the NADH nicotinamide group and may stabilize bound dinucleotide (Figure 3A). Both Arg320^F^ and His400^F^ may serve to counter-balance the negative charges of electrons on N1a and N3 clusters and to increase protein stability. The structure does not reveal specific features explaining the decreased affinity for FMN in the reduced enzyme (Holt et al., 2016). This can be attributed to minor conformational changes in the pocket upon enzyme reduction.

The Q-binding site in complex I is formed at the end of a crevice between NuoD and NuoB subunits (Baradaran et al., 2013). In *E. coli,* this wedge is formed by the 58–69 stretch of NuoB and the tip of the 220–225 loop from subunit NuoD. Both ^D^Tyr273 and ^D^His224, found in the proximity of bound decylubiquinone (Baradaran et al., 2013) are conserved in *E. coli* and point towards the quinone binding site, whereas the tip of the 218–223 loop is disordered as in most complex I structures.

### Environment and potentials of iron-sulfur clusters

At a resolution of 2.1 Å the atoms constituting the iron-sulfur clusters are resolved as independent density blobs. The conformation of side chains as well as the positions of hydrating waters in the primary and secondary interaction spheres are mostly unambiguously resolved (Figure 3). In *E. coli* complex I, cluster N1a can be reduced by NADH due to its uniquely high potential (∼ -0.3 V), differentiating it from other characterized species in which N1a cannot be reduced by NADH (Birrell et al., 2013; Zu et al., 2002). The potential of iron-sulfur clusters in proteins among other factors depends on solvent exposure, proximity of charged residues, and the number of hydrogen bonds formed between the cluster environment and sulfur atoms of clusters or coordinating cysteines (Denke et al., 1998; Fritz et al., 2002). Comparison of the chemical environment of N1a with other high-resolution structures of complex I revealed three specific differences explaining the higher potential of the N1a cluster (Table 5): (1) *E. coli*-specific ^E^Asn142 forms a hydrogen bond with S*γ* of ^E^Cys97 coordinating the N1a cluster and with N1a S1 (Figure 3B), consistent with its mutation to Met decreasing potential by 53 mV (Birrell et al., 2013). (2) In *E. coli*, water molecule W74 forms a hydrogen bond with S*γ* of ^E^Cys97. This water molecule resides in a hydrophilic cavity created by *E. coli* specific ^E^Gly140, replacing the alanine residue found in other species. (3) Because of small differences in the backbone conformation of NuoF, the backbone nitrogen of ^F^Gly97 can form a hydrogen bond with S*γ* of ^E^Cys133 in *E. coli* and *Aquifex aeolicus* but not in *Ovis aries* (Figure 3B, Table 5).

**Table 5.** Differences in the hydrogen bond network of iron-sulfur clusters in complex I structures solved at high resolution and water molecules in the immediate cluster environment. (Only the clusters for which such comparison could have been done and clusters displaying differences in the environment are listed)

| Cluster, Subunit | Organism |  |  |
| --- | --- | --- | --- |
|  | *E. coli*<br>this work | *A. aeolicus*<br>[pdb: 6hla] | *O. aries*<br>[pdb: 6zk9] |
| N3, NuoF | His400 (+)<br>Trp363(-)<br>Asn196(+) | Leu395(-)<br>Glu349(+)<br>His198(-) | Leu407(-)<br>Gln361(+)<br>Lys202(-) |
| N1b, NuoG | HOH386 |  | HOH1070 |
| N7, NuoG | HOH441<br>HOH577<br>Cys228<br>Cys231<br>Cys235<br>Cys263 |  | Asp229<br>Asp232<br>Ser236<br>Ser264<br>HOH929*<br>HOH933*<br>HOH1005* |
| N4, NuoG | Thr203(+) |  | Val205(-) |
| N5, NuoG | conserved |  |  |
| N6a, NuoI | Phe92(-) |  | Tyr109(+)<br>HOH539 |
| N6b, NuoI | Leu48(-)<br>Cys74(+)<br>Leu116(-) |  | His65(+)<br>Ala91(-)<br>Glu133(+) |
| N2, NuoB | HOH438<br>HOH211<br>Arg250<sup>D</sup><br>Ser62 |  | HOH353<br>HOH579<br>Arg85<br>dimethylated<br>Ala53 |

\* Water molecules replacing the N7 cluster

**Table 6.** Histidine residues with unassigned features extending from the imidazole ring

| Residue | Modelled atom | Comments |
| --- | --- | --- |
| <sup>E</sup>His87 | None | Interaction with <sup>E</sup>D146 |
| <sup>E</sup>His150 | HOH | Distance 2.1 Å |
| <sup>E</sup>His152 | HOH | Distance 2.4-2.6 Å |
| <sup>G</sup>His5 | HOH | Density on both sides |
| <sup>G</sup>His101 | HOH | Positive environment |
| <sup>G</sup>His123 | HOH | Distance 2.2Å |
| <sup>G</sup>His427 | HOH | Distance 2.85 Å |
| <sup>G</sup>His653 | HOH | Distance 2.5 Å very strong |
| <sup>CD</sup>His163 | HOH | Distance 2.5 Å |
| <sup>CD</sup>His507 | HOH | Distance 4.2 Å |

The environment of the other iron-sulfur clusters is mainly conserved. The differences in hydrogen donors to the clusters, cysteine sulfur atoms, and water molecules in the cluster vicinity are listed in Table 4. Clusters N3 and N2 are briefly discussed below as being the most interesting.

Cluster N3 interacts with ^F^His400, which is absent in other structurally characterized species; however, the potential of N3 is very similar between species (Leif et al., 1995; Yagi and Matsuno-Yagi, 2003). The effect of proximal His residue is likely compensated by ^F^Trp363 replacing the hydrogen bond donors (Glu or Gln) found in other species (Table 4).

The potential of cluster N2, the electron donor to quinone, varies in different species (Hirst and Roessler, 2016) notably being lower in *E. coli* compared to its mammalian analogues (-220 mV *vs.* -140 mV, respectively). However, the structure shows that the polar environment of N2 is very conserved (Figure 3C), including two water molecules, W211 and W438. Two arginines found in close proximity to the N2 cluster, Arg270^D^ and Arg250^D^, have conserved positions despite ^49kDa^Arg85 in the mammalian homologue (^D^Arg250) being dimethylated (Carroll et al., 2013). This modification prevents it from forming a hydrogen bond with ^B^Cys63, which should decrease N2 potential in the mitochondrial enzyme. Therefore, finer structural differences including those in cluster geometry, are likely responsible for differences in potential, which can likely be explained by high-resolution structure-based modeling.

### Structure of the membrane arm

The model of complete membrane arm, including the previously missing subunit NuoH (Efremov and Sazanov, 2011), was built into the density map with local resolution better than 3.5 Å at the arm center and approximately 4.0 Å at its periphery (Figure 1A, Figure 1 - figure supplement 4). An additional density belt corresponding to the lipid nanodisc is clearly visible (Figure 1A, Movie 1) around the membrane-embedded region. It is flat in the plane of the membrane with a thickness of approximately 30 Å, and closely matches hydrophobic surface of the membrane arm. The belt locally bends next to the subunit NuoL at the region where it interacts with the long amphipathic helix and is thinned next to the ^H^TMH1 (Movie 1).

The structure of the membrane arm in the lipid nanodisc is very similar to the crystal structure of the detergent-solubilized membrane arm (Efremov and Sazanov, 2011) (RMSD of 1.1 Å over 12662 atoms) (Figure 4 - figure supplement 1). The curvature of the membrane arm observed previously (Efremov and Sazanov, 2011) was unchanged in the lipid environment, and therefore, is not an artifact of crystallization or solubilization (Verkhovskaya and Bloch, 2012). Local structural differences in crystal structure include expected repositioning of ^A^TMH1 next to ^H^TMH2 (Baradaran et al., 2013), and a change in conformation of the ^M^TMH5-TMH6 loop (Figure 4 - figure supplement 1).

**Figure 4.**
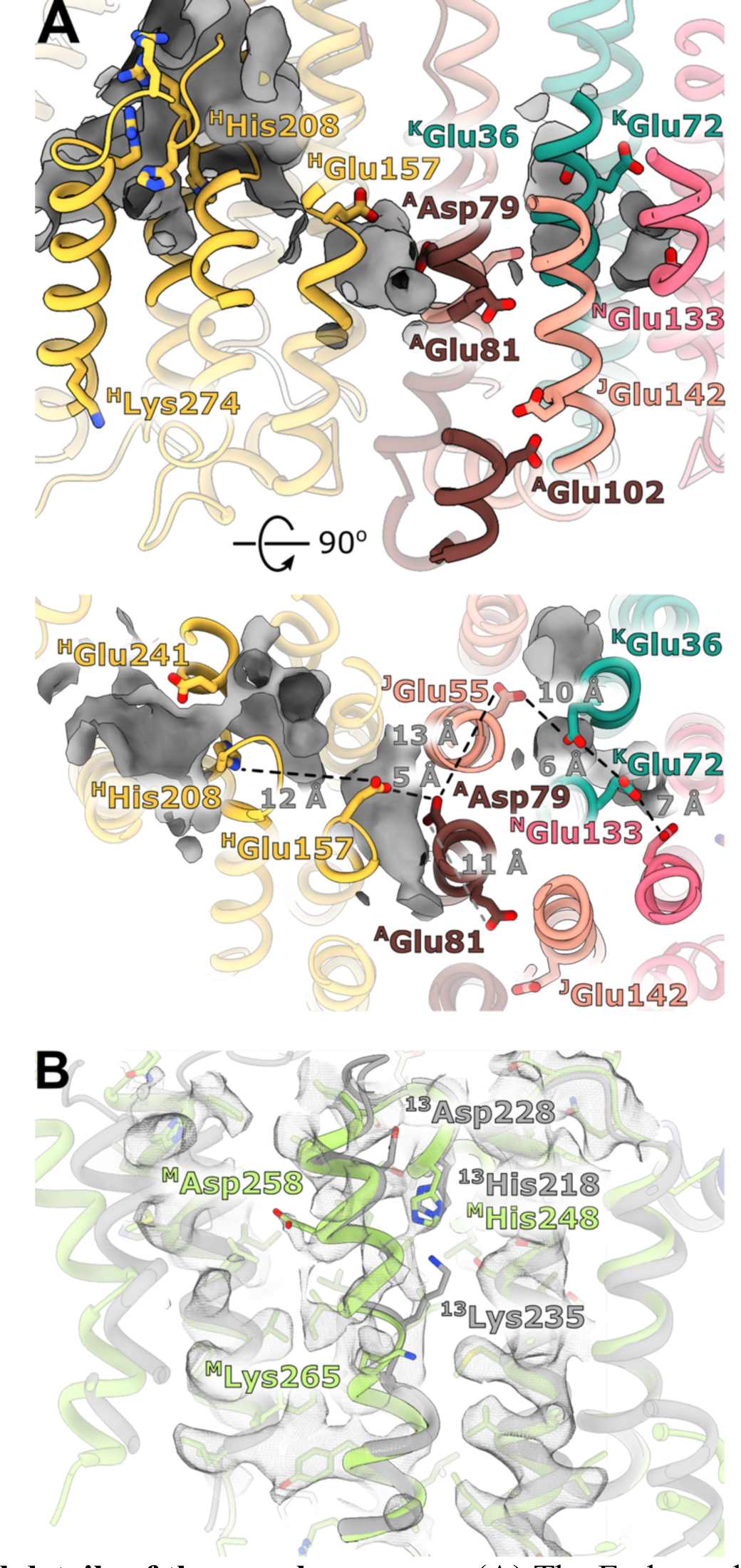
Structural details of the membrane arm. (A) The E-channel. Top: Side view, bottom: view from the cytoplasm. Charged residues between NuoH and NuoN subunits are indicated as along with the distances between them. The cavities allowing entrance of ions and water molecules are shown as grey surfaces. (B) Conformational heterogeneity within the NuoM subunit. The *E. coli* structure and amino acids are green-colored, whereas the aligned structure of *T. thermophilus* is grey. The *E. coli* membrane arm density is depicted.

The fold of subunit NuoH is similar to the structures of *T. thermophilus* and of eukaryotic complexes with one important exception. The density for the N-terminus of NuoH (residues 1-52) that includes ^H^TMH1 and a part of ^H^TMH1-TMH2 loop, is completely missing in the reconstructions of the membrane fragment and of complete complex I (Figure 1, Movie 1) suggesting that ^H^TMH1 is very mobile in the lipid nanodisc. This helix is close to the border of the nanodisc, and nanodisc belt is thinned on the cytoplasmic surface of the nanodisc at the position where the density of ^H^TMH1 disappears (Movie 1). Simultaneously there is sufficient room to accommodate the trans-membrane helix within the nanodisc.

The structures of the membrane fragment and entire complex I visualize a complete chain of charged residues connecting the Q-site with charged residues in antiporter-like subunits. (Figure 4A). We analyzed the environment of ionizable residues found within the ‘E-channel’ (Baradaran et al., 2013), a region situated between the Q-cavity and antiporter-like subunit NuoN, to evaluate the existence of a continuous proton translocation path linking the Q-cavity with the antiporter-like subunits suggested for ovine complex I (Kampjut and Sazanov, 2020). The trans-membrane region of *E. coli* NuoH contains fewer charged residues than its homologues from other structurally characterized species (Figure 4 - figure supplement 2).

Here, only *E. coli*-specific ^H^His208, separated from ^H^Glu157 by 12 Å, is found in the center of the membrane-embedded region of NuoH (Figure 4A). However, a large hydrophilic cavity stretches from the Q-site towards the center of subunit NuoH, ending next to the invariant **^H^Glu157** (hereafter, invariant residues are marked in bold). Although **^H^Glu157** is not directly linked to the cavity, DOWSER++ (Morozenko and Stuchebrukhov, 2016) placed waters linking it to the cavity, suggesting that this glutamic acid can exchange protons with the Q- cavity.

The region between NuoH and NuoN includes 6 ionizable side chains located in the middle of the membrane bilayer, 4 of which are invariant (Figure 4A,B). The distances between the residues vary from 5 Å to 12 Å which requires either displacement of the side chains or presence of water molecules to enable proton exchange between them. Analysis of cavities and potential hydration sites using DOWSER++ shows that the cluster of **^H^Glu157** and **^A^Asp79** along with the carbonyl oxygen of ^J^Gly61 forming a *π*-bulge on the ^J^TM3 (similar to the X-ray structure) (Efremov and Sazanov, 2011), indicate a hydrophilic cavity that can accommodate several water molecules, enabling proton exchange between these two residues. Carboxyl groups of a chain comprising ^J^Glu55-**^K^Glu36**-^K^Glu72-**^N^Glu133** are separated by cavities that can potentially be hydrated, enabling proton exchange between the residues. In *E. coli* complex I, residues **^A^Asp79** and ^J^Glu55/ **^K^Glu36** are separated by a distance exceeding 12 Å and a region packed with hydrophobic residues, making proton exchange between the Q- site and NuoN unlikely. **^A^Glu81,** located opposite **^A^Asp79** on ^A^TMH2, apparently does not participate in linking **^A^Asp79** with **^N^Glu133.** However, it faces hydrophilic environment of ^J^Ser145, *E. coli*-specific ^J^Glu142, and ^A^Glu102, potentially linking it to the periplasmic surface (Figure 4A). Our analysis indicated that in *E. coli,* the E-channel is less pronounced than in *T. thermophilus* and that no continuous proton path exists between the Q-site and NuoN.

Curiously, ^H^Lys274 almost universally conserved in complex I and related hydrogenases is found in the ^H^TMH7 off the main pathways proposed for proton translocation. In our structure, its ammonium group is oriented towards the periplasm (Figure 4A); however, the length and flexibility of the side chain would allow it to reach the center of the membrane upon structural rearrangement.

The cytoplasmic half of ^J^TMH3 was found to assume two alternative conformations in eukaryotic complex I (Agip et al., 2018; Kampjut and Sazanov, 2020). In *E. coli* complex I, the density in this region is very well-resolved, suggesting the absence of alternative conformations. A peculiar feature is observed in the density of subunit NuoM instead.

The density of the cytoplasmic half of ^M^TM8 is poor and fragmented between residues 255 and 265, indicating the existence of multiple conformations (Figure 4C, Movie 2). This region is buried in the middle of NuoM and the density of surrounding helices is very well-resolved indicating the local character of the disorder. This region spans the invariant **Lys265** including the *π*-bulge, and found in some bacteria ^M^Asp258. Interestingly, though the helix structure in mammalian complex I is similar to that in *E. coli* (the equivalent Asp is missing), the corresponding region in *T. thermophilus* differs significantly (Figure 4C). The cytoplasmic region of ^13^TM8 is rotated by 2 residues. This is achieved by extending TMH8 at *T. thermophilus* **^13^Lys 235** (equivalent to *E. coli* **^M^Lys265**). In *E. coli*, **^M^Lys265** is buried in the center of the second structural repeat whereas in *T. themophilus*, **^13^Lys 235** is positioned at the interface between the structural repeats facing **^13^His218** (in *E. coli* ^M^His248). This rotation places ^13^Asp228 (equivalent to *E. coli* ^M^Asp258) pointing towards central axis of the first structural repeat (TM4-8) and exposed to the cytoplasm, whereas in *E. coli*, it is buried on the central axis of the second structural repeat (TM9-TM13). Thus, higher mobility of the helical fragment situated at a critical position at the interface of symmetry-related modules may indicate *π*-bulge-enabled helical rotation with a possible role in proton translocation.

### The peripheral-membrane arm interface

The interface between membrane and peripheral arms presents an important element of the complex that mediates the coupling of ubiquinone reduction above the cytoplasmic surface to proton translocation across the membrane. The interface between peripheral and membrane arms is primarily formed through interaction between subunits NuoB and NuoD of the peripheral arm with the cytoplasmic surface of NuoH and the TMH1-TMH2 loop of subunit NuoA in the membrane domain. The residues involved in the direct interaction between the arms and the interface structure are highly conserved (Figure 5 - figure supplement 1) between all complex I and related membrane-bound hydrogenases (Baradaran et al., 2013; Grba and Hirst, 2020; Kampjut and Sazanov, 2020; Yu et al., 2020; 2018).

**Figure 5.**
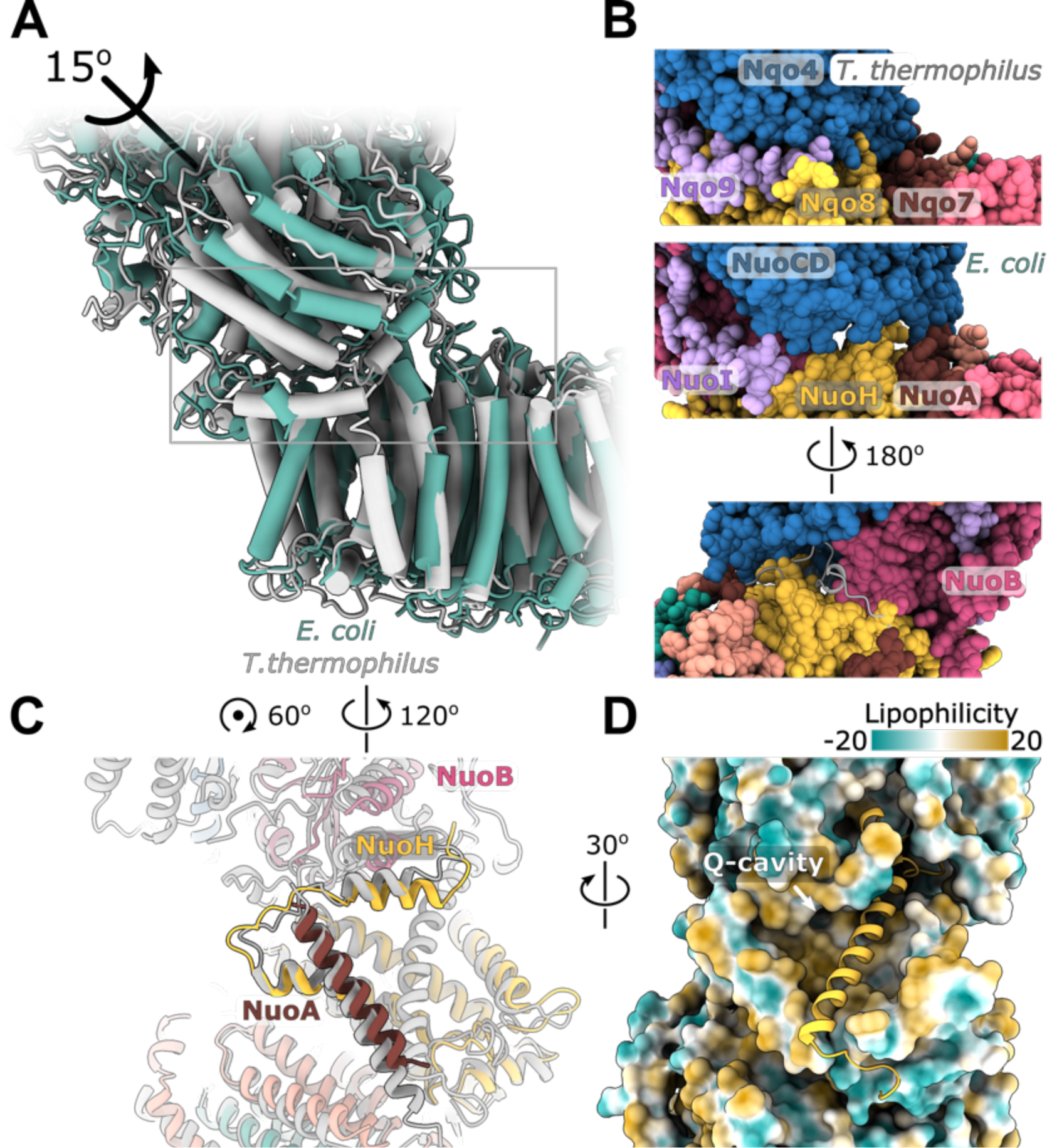
Interface between the peripheral and membrane arms. (A) Comparison of the interface between *E. coli* (green) and *T. thermophilus* (PDB ID:4HEA, grey) complex I. Structures were aligned on the subunit NuoH/Nqo8. The rotation axis of the subunits NuoB/D module relative to Nqo6/4 is indicated. (B) Interfacial contacts between the peripheral and membrane arms in *T. thermophilus* (upper panel) and *E. coli* (middle and bottom panel). A gap in the subunit interface is apparent in the absence of the conserved ^A^TMH1-2 loop fragment. The corresponding loop from *T. thermophilus* is shown in grey in the cartoon representation (bottom panel). (C) Differences in the structures of NuoH and NuoA subunits between *E. coli* (colour coded as in Figure 1) and *T. thermophilus* (grey). (D) View from the membrane on the entrance to the Q-cavity. Homology model of ^H^TM1, absent in the *E. coli* structure, is shown in the cartoon representation. The protein surface is coloured by lipophilicity.

Local resolution (Figure 1 - figure supplement 4) and B-factors (Figure 5 - figure supplement 2) of the *E. coli* peripheral arm show that the membrane-facing surface (subunits NuoD and NuoB) including the residues lining the Q-cavity are significantly less ordered than the remaining subcomplex. The mobility in the interfacial region of the arm is very similar in the reconstructions of the peripheral arm dissociated from and complexed with the membrane arm. However, several interfacial regions of subunits NuoB, NuoD, and NuoI become more ordered upon complex formation and their density can be observed in the reconstruction of isolated conformations of the entire complex I (Figure 5 - figure supplement 2, Table 2). Similar to complex I from *T. thermophilus* (Baradaran et al., 2013), there are no specific conformational changes at the interface upon association of the arms. These results suggest that the interfacial region of the peripheral arm is inherently flexible and likely responsible, at least in part, for the high relative mobility of the arms.

Structure of the surface of subunit NuoH and relative arrangement of subunits NuoB and NuoD in *E. coli* complex I is similar to that of complex I from other species (RMSD 1.2 Å over 324 C*α* atoms with *T. thermophilus* complex I). However, their relative positions differ. Thus, in *E. coli* complex I, NuoB and NuoD are rotated around an axis passing through the center of NuoH and the interface between NuoF and NuoG anticlockwise when observed from the top of peripheral arm at approximately 15 degrees (Figure 5A). This results in the shift of NuoD interfacial regions with an amplitude exceeding 10 Å and the separation of NuoD from NuoH, which reduces the interaction between the four-helical bundle domain of NuoD with NuoH (Figure 5B). The highly conserved fragment of the ^A^TMH1-TMH2 loop (residues 46-53), that forms a plug between subunits NuoD and NuoB (Figure 5 - figure supplement 1B) and interacts with the ^D^221-228 loop containing ubiquinone-coordinating His224, is also disordered in our structure (Figure 5B).

On the opposite side of the interface, structural rearrangements include a 7-degree tilt of ^A^TMH1 that becomes more perpendicular to the membrane plane and approximately 15- degree rotation of the amphipathic helix in the loop connecting ^H^TMH1-TMH2, residues 57– 68, in the direction of ^H^TMH1 and towards the membrane center (Figure 5C). Analysis of the three full conformations identifies this helix as the main membrane arm element that performs rearrangement together with the cytoplasmic domain. Its rotation reduces the opening to the Q-cavity (Figure 5D). Homology modeling indicates that the observed rearrangements are still compatible with ^H^TMH1 occupying its expected position without any steric clashes (Figure 5D) suggesting that ^H^TMH1 is highly mobile in the lipid environment rather than being absent from its expected position.

Rotation of the NuoB/NuoD subunits module creates multiple openings on the interface between the arms (Figure 5B). The size of the openings is compatible with the diffusion of water molecules and likely, of protons from the outer space towards the Q-cavity. Such openings suggest that ubiquinone bound to the Q-site can receive protons directly from the solvent.

## Discussion

### Structural features of *E. coli* complex I

*E. coli* complex I is composed of the smallest number of subunits among all complex I structures characterized so far. Yet it still evolved a strategy to stabilize peripheral arm assembly without involving additional subunits. The interactions between subunits are stabilized by extended C-termini and a large G-loop (Figure 2), in turn stabilized by the Ca^2+^ ion, which is known to modulate the complex stability (Sazanov, 2003). This indicates an evolutionary pressure on maintaining the peripheral arm integrity, which was ‘solved’ in a species-dependent manner.

There is no apparent continuous proton translocation path between the Q-cavity and subunit NuoN in *E. coli* complex I. Further, there are no indications for the existence of different conformations in the cytoplasmic half of ^J^TMH3 observed in mammalian complex I (Agip et al., 2018; Kampjut and Sazanov, 2020) attributed to deactive-active transition (Agip et al., 2018) or more recently, to different catalytic intermediates (Kampjut and Sazanov, 2020).

This suggests that these states are either suppressed in the resting state of the bacterial complex or do not occur at all. Conversely, ^M^TMH8 displays localized disorder next to the *π*- bulge, indicating involvement of this helix in the structural rearrangements associated with proton translocation, and to our knowledge, represents the first indication of specific conformational changes in antiporter-like subunits.

*E. coli* complex I is known to be a dynamic complex (Morgan and Sazanov, 2008; Sazanov, 2003). Our cryo-EM reconstructions reveal the reasons for its high flexibility. The peripheral and membrane arms are mainly rigid, whereas the connection between arms is flexible (Figure 1C, Figure 1 – figure supplement 5). Two reasons can be identified for this: 1) high mobility of the interfacial regions of subunits NuoB and NuoD (Figure 1 - figure supplement 4, Figure 5 - figure supplement 2) and 2) the 15-degree rotation of the interfacial subunits NuoB and NuoD relative to NuoH, observed uniquely in *E. coli* (Figure 5A). The rotation disrupts many conserved complimentary interactions between the arms and renders the interface porous such that the Q-cavity is exposed to the solvent. This is different from all the other known structures of complex I and evolutionarily related complexes in which the interface is solvent-inaccessible. Therefore, we interpreted the observed conformation of *E. coli* complex I as an uncoupled state. Our preparation of the complex is competent in ubiquinone reduction (Figure 1 - figure supplement 2); however, unless conformational changes sealing the Q-cavity occur during the catalytic cycle, ubiquinone reduction by NADH in the present conformation is expected to occur without proton translocation.

The reasons for this difference in interface conformation with other structurally characterized complexes are not clear. It may represent a resting state described in *E. coli* complex I (Belevich et al., 2017), which like in eukaryotes (Babot et al., 2014), is characterized by lower catalytic activity and is activated by NADH:ubiquinone oxidoreduction cycles. However, reactive states in eukaryotic complex I are associated with local conformational changes involving loop rearrangement (Agip et al., 2018; Parey et al., 2018), rather than displacement of the complete domains observed in *E. coli* complex I. Other reasons for the observed rotation of arms may include the higher concentration of divalent ions used in our sample that weakens multiple salt bridges linking the arms, or displacement of the amphipathic termini of subunits NuoB and NuoI due to the limited size of the nanodisc, resulting in weakened interaction between the arms.

The absence of density for ^H^TMH1 is another unique feature of *E. coli* complex I. It likely reflects the higher dynamics of this helix in lipid environment. Possibly, tilted ^A^TMH1 (Figure 5C) displaces the periplasmic end of ^H^TMH1 disrupting its interaction with the TMH2-3 loop and TMH6 of NuoH, rendering this helix dynamic. It may also have a functional role because helix displacement facilitates the otherwise too narrow access to the Q-cavity for ubiquinone (Baradaran et al., 2013). Further insights into the role of ^H^TMH1 might be obtained once conditions stabilizing the coupled complex conformation are identified.

### Revised coupling mechanism

Based on structural features of *E. coli* complex I and the wealth of available structural information, we would like to propose a coupling mechanism that differs from those suggested previously (Figure 6), which is simple, compatible with the microscopic reversibility principle (Onsager et al., 1996), has evolutionary meaning, and is applicable to the entire class of evolutionarily related complexes.

**Figure 6.**
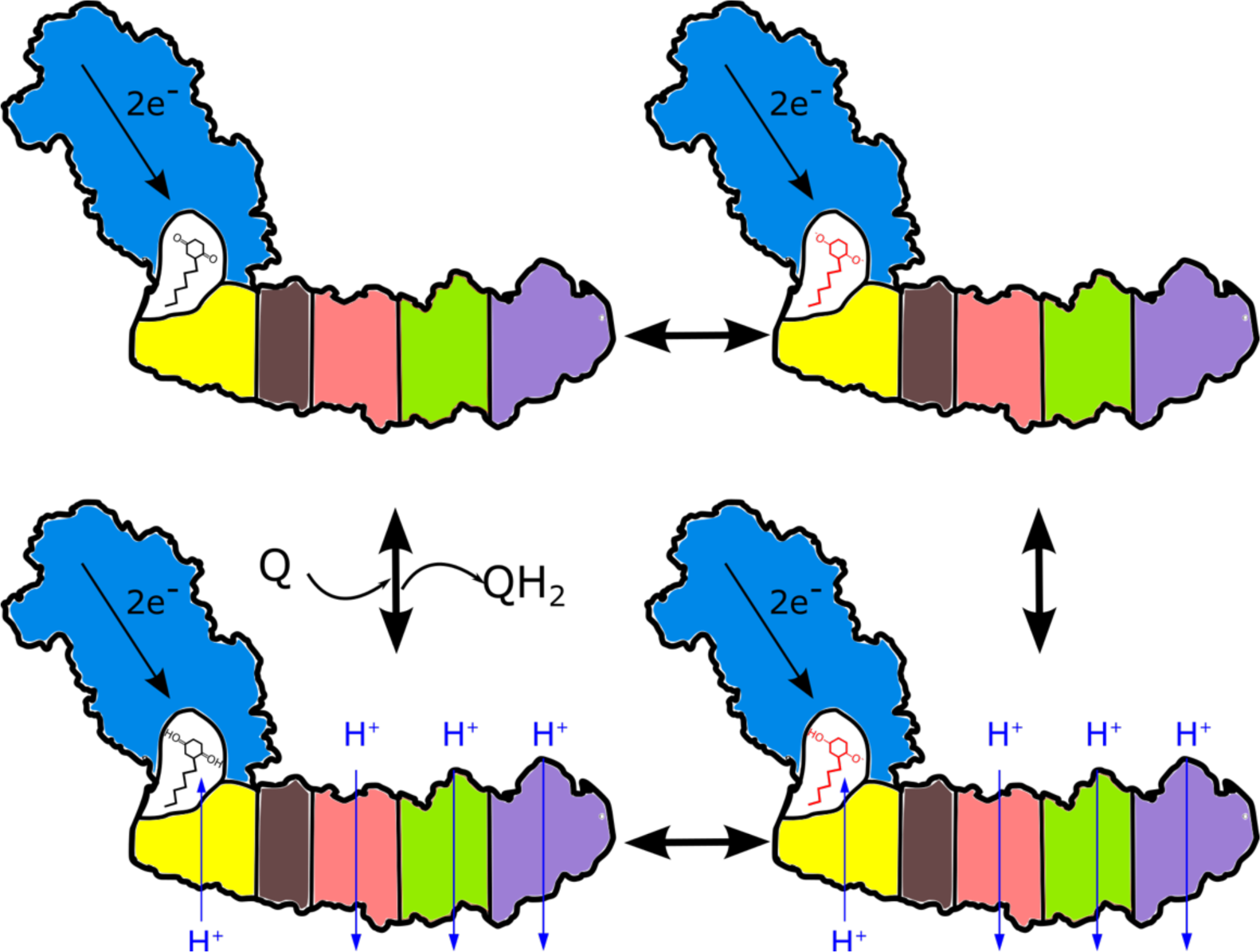
Proposed mechanism of coupling in respiratory complex I. Ubiquinone reduction decreases the proton potential in the Q-cavity, generating/enhancing electrochemical potential between the Q-cavity and periplasmic space. It is subsequently neutralized by protons translocated through NuoH from the periplasm into the Q-cavity and this translocation is coupled with the reversible translocation of three protons into the periplasm. Colour coding of schematic subunits in the membrane arm is similar to that described in Figure 1, negatively charged states of ubiquinone are shown in red.

We propose that the key to the coupling is the formation of a cavity isolated from external protons and accessible to ubiquinone such that ubiquinone can exchange electrons with the N2 cluster. The necessity of having a tightly coupled cavity explains the high conservation of the subunit interface. Notably, as the ubiquinone entrance is situated in the hydrophobic region of the bilayer, these two requirements do not contradict each other.

The potential of benzoquinone-hydroquinone couple depends on the pH (Chambers, 1988), similar to that of any redox reaction involving protons, and has been shown experimentally to decrease by over 400 mV to below -300 mV upon pH change from 2 to 10 (Lemmer et al., 2011). Rough estimations indicate that addition or extraction of a single proton from a cavity with the characteristic dimensions of the Q-cavity, alters the activity of protons within the cavity by hundreds of millivolts. Thus, the redox potential of ubiquinone bound within the cavity enclosed from the environment will be strongly modulated by the extraction/addition of single protons from/to the cavity. *Vice versa*, reduction or oxidation of ubiquinone/ubiquinol is equivalent to adding/removing proton binding groups to/from the Q-cavity. Thus, ubiquinone serves as a transformer that converts the energy of electrons to the chemical potential of protons in a fully reversible manner. In the coupled complex during the forward cycle, ubiquinone reduction decreases proton activity in the cavity, which is rectified by protons entering the cavity and performing work. Questions of how the protons perform the work and where they come from are thus critical to formulate the coupling mechanism.

Multiple proton pathways have been suggested (Baradaran et al., 2013; Efremov and Sazanov, 2012; Kampjut and Sazanov, 2020; Verkhovskaya and Bloch, 2012; Yu et al., 2020; 2018). However, they all end up on intracellular/matrix side of the membrane, which makes it difficult to explain the energy conversion mechanism. Instead, we propose that the protons re- protonate ubiquinone through NuoH and/or adjacent trans-membrane subunits from the periplasmic/inter membrane side of the membrane as shown in Figure 6. The fold of subunit NuoH contains a set of 5 TMH with a helical arrangement similar to the symmetric module in antiporter-like subunits including broken TMH (Baradaran et al., 2013) and invariant ^H^Glu157 in a position similar to the invariant ^M^Glu144, suggesting that it can translocate protons. Proton transport through NuoH is coupled to the transport of three protons by three antiporter-like subunits in the opposite direction. This coupling must involve both the interaction of ionizable residues in the middle of the membrane (Baradaran et al., 2013; Efremov and Sazanov, 2011) as well as conformational changes, and likely proceeds through a classical alternating access mechanism (Jardetzky, 1966). Thus, the entire membrane module functions as a reversible proton antiporter with the stoichiometry of 1H^+^in/3H^+^out. Four protons are translocated outside in two pumping cycles per one reduced ubiquinone molecule (Figure 6). In this mechanism, ubiquinone reduction creates a local enhanced membrane potential on the NuoH subunit between the Q-cavity and periplasmic space. Moreover, only the equilibrium potential of NADH and ubiquinone as well as the transmembrane potential are important for the directionality of the reaction and energy balance as expected for a molecular machine (Astumian et al., 2016).

Under equilibrium, the potential of ubiquinone in the Q-cavity is equilibrated with the potential of NADH, and of protons in the Q-cavity, which results in trans-membrane potential-dependent semiquinone species as observed by EPR spectroscopy in tightly coupled submitochondrial particles (Yano et al., 2005) .

The proposed mechanism is applicable to all the complexes that are evolutionary related to complex I (Efremov and Sazanov, 2012; Yu et al., 2020; 2018). In all of these, the cavity formed between the peripheral arm and NuoH subunit is sealed. The peripheral arm-NuoH complex is undoubtably one of the stand-alone evolutionary modules. This is supported by the differences in its position between complex I and membrane-bound hydrogenases (Yu et al., 2020; 2018), and its susceptibility to dissociation from the membrane arm in *E. coli* (Baranova et al., 2007; Efremov and Sazanov, 2011) as expected for a late evolutionary addition (Levy et al., 2008). The initial association of the hydrogen-evolving module with an antiporter may have had an evolutionary advantage with the proton-translocating module serving as a source of protons (Yu et al., 2018), biasing H2 evolution towards the reaction product or to enhance Na^+^ extraction from the cells (Boyd et al., 2014).

Complex uncoupling is achieved by opening the Q-cavity to the solvent, consistent with elegant experiments by Cabrera-Orefice et.al (Cabrera-Orefice et al., 2018) in which the locking conserved the ^A^TMH1-2 plug with the cysteine bridge reversibly uncoupled the enzyme. Close examination of the crosslinked structure indicates that crosslinking fixes the plug conformation in a way that the Q-cavity is accessible to the solvent.

The exact proton translocation mechanism within the antiporter-like module is unknown and requires further experimental and computational investigation. Here, we can only speculate that given the high conservation of ^H^Glu157, it plays an important role in the coupling and likely changes its protonation state during the pumping cycle. Thus, it can influence the pKa of neighboring ionizable residues. In membrane-bound hydrogenase (MBH), an equivalent ^M^Glu141 is separated from the closest ionizable ^H^Lys409 by distance of 20 Å, which in a hydrophobic environment with a dielectric constant of 10, allow them to mutually modulate the pKa of each other by approximately 1 pH unit, similar to the free energy conserved upon ferredoxin oxidation by the protein complex. This distance is reduced to around 13 Å in membrane-bound sulfane sulfur reductase (MBS) and to around 6 Å in complex I, consistent with the proportionally higher free energy of catalyzed reactions (Yu et al., 2020; 2018).

The proposed coupling mechanism also suggests how the different conformational states associated with proton translocation might be trapped. A pH jump applied to purified coupled complex I will create a difference in potential between the Q-cavity and periplasmic surface, which depending on the direction of the jump, may trap different equilibrium conformations of this molecular machine.

## Materials and Methods

### Generation of an *E. coli* strain expressing Twin-Strep-tagged respiratory complex I

The native *nuo* operon encoding the 13 subunits of respiratory complex I (NuoA-N) was engineered with a Twin-Strep-tag (WSHPQFEKGGGSGGGSGGSAWSHPQFEK, IBA GmbH) at the N-terminus of NuoF using CRISPR-Cas9-enabled recombineering(Jiang et al., 2015). The DNA sequence encoding the C-terminal region of NuoE and N-terminus of NuoF was retrieved from GenBank (Acc. No. NC_012971.2 region 2288438 – 2289174). The tag-coding sequence followed by a TEV protease recognition site (Tropea et al., 2009) was appended upstream of the NuoF N-terminus and was codon-optimized, together with the 2288766–2288807 region of the genomic fragment. Such designed, linear DNA knock-in cassette was synthesized (GenScript). The vectors pCas and pTargetF were gifts from Sheng Yang (Addgene plasmids #62225 and #62226). The N20 sequence (GGTCAGCGGATGCGTTTCGG) was introduced into pTargetF by inverse PCR. Genomic engineering was performed according as described by Jiang *et al* (Jiang et al., 2015). Briefly, pCas vector was transformed into the chemically competent *E. coli* BL21AI strain (Thermo Fisher Scientific Inc.). The transformants were grown in shake-flask culture at 30°C in Lysogeny Broth (LB) medium containing 25 μg mL^-1^ (w/v) kanamycin monosulfate and 10 mM L-arabinose. Upon reaching OD600 0.5, the bacteria were rendered electrocompetent and were co-electroporated with the linear DNA cassette and the mutated pTargetF vector. The transformants were selected on LB-agar plates supplemented with 25 μg mL^-1^ (w/v) kanamycin and 50 μg mL^-1^ (w/v) streptomycin, or 50 μg mL^-1^ (w/v) spectinomycin. The positives, identified by colony PCR and DNA sequencing, were cured of the plasmids as described previously (Jiang et al., 2015). We further refer to the modified strain as *E. coli* BL21FS (NuoF-Strep).

### Expression and purification of respiratory complex I

*E. coli* BL21FS was cultivated in LB medium for 48 hours at 37°C in a microaerobic environment. The cells were harvested by centrifugation and the membrane fraction was isolated as described by Sazanov *et al* (Sazanov, 2003). All subsequent steps were performed at 4°C. The homogenate was solubilized in 2% (w/v) n-Dodecyl β-D-maltoside (DDM, Anatrace) for 2 hours while stirring, after which the non-solubilized fraction was removed by ultracentrifugation at 225 000 × *g* for 1 hour. The supernatant was adjusted to 200 mM NaCl and loaded on a 5 mL Strep-Tactin® Superflow® high capacity column (IBA GmbH). After washing with 25 column volumes (CV) of buffer A (50 mM Bis-tris pH 6, 2 mM CaCl2, 200 mM NaCl, 0.04% (w/v) DDM, 10% (v/v) sucrose, 0.003% (w/v) *E. coli* polar lipid extract (Avanti Polar Lipids, EPL), 0.2 mM PMSF), complex I was eluted with 2 CV of buffer B (buffer A containing 5 mM D-desthiobiotin (IBA GmbH). The purity of the eluted protein was assessed by SDS-PAGE and activity assays (Figure 1 - figure supplement 1). The purified complex I was concentrated using an Amicon Ultra-4 100K centrifugal filter (Merck) to 0.5 mg mL^-1^ (w/v), fast-frozen in liquid nitrogen and stored at -80°C.

### Reconstitution of respiratory complex I into lipid nanodiscs

The membrane scaffold protein MSP2N2 was expressed and purified following a published protocol (Grinkova et al., 2010). Purified complex I at concentration 520 nM was mixed with 10.4 µM MSP2N2 (1:20 protein:MSP molar ratio) and incubated for 1 hour at 4°C.

Subsequently, the detergent was removed by adding 0.5 g mL^-1^ (w/v) Bio-Beads (Bio-Rad) overnight at 4°C. The reconstituted protein was further purified on the Superose 6 Increase 10/300 GL column (GE Healthcare) equilibrated in a buffer comprising 20 mM Bis-Tris pH 6.8, 200 mM NaCl and 2 mM CaCl2. The protein-containing fractions were pooled and concentrated to 0.1–0.2 mg mL^-1^ (w/v) using Amicon Ultra-0.5 100K centrifugal concentrators.

### Activity assays

NADH:ferricyanide (FeCy) and NADH:ubiquinone-1 (Q1) activities were measured as described previously (Sazanov, 2003). For the assays, 3 nM of detergent-solubilized or nanodisc-reconstituted complex I and either 1 mM FeCy (Sigma Aldrich BVBA) or 100 μM Q1 (Sigma Aldrich BVBA) were added to the assay buffer (10 mM Bis-Tris pH 6.8, 200 mM NaCl, 10 mM CaCl2) in a stirred quartz cuvette at 30°C. The reaction was initiated by adding 100 μM NADH (Carl-Roth GmbH) and followed as reduction in absorbance at 340 nm using a Varian Cary 300 UV-Vis spectrophotometer (Agilent Technologies, Inc). During the inhibition assay, complex I was incubated with 20 µM Piericidin A (Cayman Chemical) for 5 min in the assay buffer at 30°C prior to Q1 addition.

### Mass photometry

The composition of the protein preparation was assessed using mass photometry on a Refeyn OneMP instrument (Refeyn Ltd.), which was calibrated using an unstained native protein ladder (NativeMark™ Unstained Protein Standard A, Thermo Fisher Scientific Inc.). Measurements were performed on the reconstituted complex I at a concentration of 0.015 mg ml^-1^ using AcquireMP 2.2.0 software and were analyzed using the DiscoverMP 2.2.0 package.

### Preparation of cryo-EM samples

The cryo-EM samples were prepared using a CP3 cryoplunge (Gatan). Quantifoil R0.6/1 Cu300 holey carbon grids were cleaned with chloroform, acetone, and isopropanol as described by Passmore *et al* (Passmore and Russo, 2016) . The grids were glow discharged in the ELMO glow discharge system (Corduan Technologies) from both sides for 2 min at 11 mA and 0.28 mbar. Four microliters of the reconstituted protein solution at 0.15 mg ml^-1^ concentration were applied on a grid and blotted from both sides for 2.2 s with Whatman No. 3 filter paper at 97 % relative humidity. The grid was then plunge-frozen in liquid ethane at - 176°C and stored in liquid nitrogen.

### Cryo-EM data collection

Cryo-EM images were collected on a JEOL CryoARM 300 microscope equipped with an in- column *Ω* energy filter (Fislage et al., 2020) at 300 kV, automatically using SerialEM 3.0.8 (Mastronarde, 2005) at a nominal magnification of 60,000 and the corresponding calibrated pixel size of 0.771 Å. Five images per single stage position were collected using a cross pattern with 3 holes along each axis (Efremov & Stroobants, 2021 in press). The 3 s exposures were dose-fractionated into 61 frames with an electron dose of 1.06 e- Å^-2^ per frame. The energy filter slit was set to 20 eV width. In total, 9122 zero-loss micrographs were recorded with the defocus varying between -0.9 and -2.2 µm (Table 1).

### EM image processing

The dose-fractionated movies were motion-corrected using MotionCor2 (Zheng et al., 2017) in the patch mode. The Contrast Transfer Function (CTF) parameters were estimated using CTFFIND-4.1 (Rohou and Grigorieff, 2015). 40 micrographs of various defoci were selected, manually picked, and used to train the neural network of crYOLO 1.4 (Wagner et al., 2019). After training, 1,256,734 particles were picked automatically from the complete dataset, extracted in RELION 3.0 (Zivanov et al., 2018), and imported into cryoSPARC 2.11^11^. Following 2D classification, six initial models were generated, among which one corresponded to the cytoplasmic arm-only and another corresponded to the complete complex I. Using hetero-refinement, 441,265 and 525,680 particles were assigned to the cytoplasmic arm and complete complex, respectively. Further processing was performed in RELION 3.1(Zivanov et al., 2020). After per-particle CTF estimation and Bayesian polishing, 3D auto- refinement of the complete complex produced a map at an average resolution of 3.4 Å (Figure 1 - figure supplement 3). However, the map was very heterogeneous with the peripheral arm resolved at 3.0–3.6 Å whereas the membrane arm was resolved at over 10 Å.

To address this heterogeneity, both arms were refined independently using multi-body refinement (Nakane et al., 2018) (Figure 1 - figure supplement 3) and the peripheral domain signal was subtracted. After two rounds of 3D classification applied to the membrane domain and nanodisc signal subtraction, a subset of 48,745 particles was 3D refined to an average resolution of 3.6 Å. However, the density map was anisotropic. To improve the reconstruction, the original stack of 525,680 particles was refined against the masked cytoplasmic arm, followed by subtraction of the signal from the peripheral arm. Next, membrane arm map obtained above was filtered to 9 Å and used as an initial model for the 3D refinement of all resulting membrane arm particles. Next, to prevent model bias, the refined map was low-pass filtered to 20 Å and used in the subsequent 3D classification with 10 classes, *τ* of 12 and 24° local angular search range and 1.8° angular step. The best class (110,258 particles and 8 Å resolution) was auto-refined using the starting model low-pass filtered to 15 Å, which produced the reconstruction to a resolution of 4.4 Å. Next, the nanodisc density was subtracted, which further improved the resolution to 3.9 Å. Following 3D classification without alignment with *τ* of 40, 8 classes, and resolution in the E-step limited to 4 Å, a subset of 37,441 particles was identified, which after auto refinement, produced a density map at an average resolution of 3.9 Å with better resolved peripheral regions. Finally, density modification with the resolve_cryo_em tool available in PHENIX 1.18.2 (Terwilliger et al., 2020) improved the resolution to 3.7 Å (Figure 1 - figure supplement 3,4).

After multibody refinement of the arms described above, peripheral arm particles with the subtracted membrane arm were 3D classified into 12 classes without alignment using *τ* of 40 and resolution of the expectation step limited to 4 Å. The best class contained 134,976 particles and was further refined to 2.9 Å resolution.

A subset of 166,580 particles was selected after a similar 3D classification procedure that was applied to the 441,265 particles of dissociated peripheral arm particles. It was further cleaned from the remaining particles of the complete complex I by 2D classification, resulting in a subset of 151,357 particles that produced a density map to a resolution of 3.0 Å. As the reconstructions of the dissociated and membrane arm-subtracted cytoplasmic arms were virtually identical, both stacks were combined. After two cycles of per-particle CTF refinement, aberration corrections, and Bayesian particle polishing in RELION 3.1, the resolution improved to 2.4 Å. Consecutive density modification in PHENIX further improved the resolution to 2.1 Å (Figure 1 - figure supplement 3,4, Table 1).

To resolve the conformation of entire complex I, a stack of 525,680 particles was aligned to the peripheral arm using auto-refinement with a mask around the peripheral arm in RELION Next, 3D classification without alignment into 30 classes with resolution of the expectation step limited to 20 Å and *τ* of 4 was performed, followed by auto-refinement of each resulting class, which produced maps to a resolution in the range of 9-20 Å (some of the classes are shown in Figure 1 - figure supplement 5).

Three high-resolution conformations of complete complex I were obtained as follows.

Conformation 1 was resolved by applying the 3D classification into 15 classes, *τ* of 6, a 24° local angular search range, and 1.8° sampling interval to the subset of 110,258 particles that produced the 3.9 Å reconstruction of the membrane arm (see above). The best class consisted of 23,445 particles that were refined to a resolution of 3.9 Å.

Conformations 2 and 3, were identified by applying 3D classification without image alignment into 12 classes with *τ* of 40 and resolution of the expectation step limited to 4 Å, to the stack of 525,680 intact complex I particles. Two of the best classes, consisting of 21,620 and 21,234 particles were refined to 4.6 Å and 4.5 Å, respectively. Following density-modification in PHENIX, the resolution of the maps was improved to 3.3, 3.8, and 3.7 Å, for conformations 1, 2, and 3, respectively (Figure 1 - figure supplement 4C, Table 1).

### Model building

Peripheral arm subunits constituting NuoB, CD, E, F, G, and I were first homology modelled in the SWISS-MODEL server (Waterhouse et al., 2018) based on the structure of *T. thermophilus* (PDB ID:4HEA (Baradaran et al., 2013)) and were rigid-body fitted into the density map in UCSF Chimera 1.13.1. Following manual rebuilding in Coot 0.9 (Casañal et al., 2020), the model was subjected to real-space refinement against the final 2.1 Å map of the cytoplasmic arm in PHENIX 1.18.2 using the default parameters. Secondary structure restrains were applied only to the interfacial region resolved at a lower resolution. The value of the nonbonded_weight parameter was optimized. Water molecules were added to the map and validated using the “Check/delete waters” tool in Coot 0.9 . Molecular dynamics-based model idealization was conducted in ISOLDE 1.0b5 (Croll, 2018), followed by several iterations of real-space refinement without atomic displacement parameter (ADP) restraints and manual rebuilding in Coot 0.9.

For the membrane domain, the previously obtained *E. coli* model (PDB ID: 3RKO) was real-space-refined in PHENIX. The missing NuoH subunit was homology-modelled using the *T. thermophilus* structure (PDB ID: 4HEA) in Coot 0.9. The final model was obtained after several rounds of manual rebuilding and real-space refinement using standard parameters with Ramachandran restrains, secondary-structure restrains applied to the NuoL TMH9-13, without ADP restrains, and with the optimized nonbonded_weight parameter. To generate the model of the complete complex I, the separate peripheral and membrane arm structures were combined and the missing parts at the interface (Table 2) were built manually. As the density of NuoL and NuoM was very poor in all the resolved full conformations, these subunits were subjected to rigid-body refinement in PHENIX, whereas the others were subjected to real-space refinement with minimization_global, local_grid_search, morphing, and ADP refinement. Ramachandran, ADP, and secondary-structure restrains were used. After manual rebuilding in Coot, real-space refinement of the full complex was performed with standard parameters and restrains. The models were validated in MolProbity (Williams et al., 2018) .

Structural conservation was evaluated using the ConSurf server (Ashkenazy et al., 2016). The figures and movies were generated in UCSF ChimeraX version 1.1. (Goddard et al., 2018) and PyMOL (The PyMOL Molecular Graphics System, Version 2.4.1 Schrödinger, LLC).

## Acknowledgements

We are indebted to Henri De Greve for help with establishing CRISPR-Cas9 for *E. coli* complex I. We are thankful to Dr. Adam Schröfel and Dr. Marcus Fislage for providing support during cryo-EM data collection, to Annelore Stroobants for technical assistance and to Lukasz Milewski for assistance in data processing. We kindly thank VIB Tech Watch fund for facilitating access to the Refeyn instrument. We would like to acknowledge the funding provided by Vlaams Instituut voor Biotechnologie, Fonds Wetenschappelijk Onderzoek (grant Nos. G0H5916N, G.0266.15N) and by the European Research Council.

## Data availability

Cryo-EM density maps and atomic models are deposited into the PDB and EMDB databases with the following accession codes: cytoplasmic domain (PDB ID: 7NZ1, EMD-12661), membrane domain (PDB ID: 7NYH, EMD-12652), entire complex conformation 1 (PDB ID: 7NYR, EMD-12653), conformation 2 (PDB ID: 7NYU, EMD-12654), conformation 3 (PDB ID: 7NYV, EMD-12655).

The following data sets were generated: Kolata P, Efremov RG (2021). Electron Microscopy Data Bank ID EMD-12661. Respiratory complex I from Escherichia coli - focused refinement of cytoplasmic arm. https://www.ebi.ac.uk/pdbe/entry/emdb/EMD-12661 Kolata P, Efremov RG (2021). Electron Microscopy Data Bank ID EMD-12652. Respiratory complex I from Escherichia coli - focused refinement of membrane arm. https://www.ebi.ac.uk/pdbe/entry/emdb/EMD-12652 Kolata P, Efremov RG (2021). Electron Microscopy Data Bank ID EMD-12653. Respiratory complex I from Escherichia coli - conformation 1. https://www.ebi.ac.uk/pdbe/entry/emdb/EMD-12653

Kolata P, Efremov RG (2021). Electron Microscopy Data Bank ID EMD-12654. Respiratory complex I from Escherichia coli - conformation 2. https://www.ebi.ac.uk/pdbe/entry/emdb/EMD-12654

Kolata P, Efremov RG (2021). Electron Microscopy Data Bank ID EMD-12655. Respiratory complex I from Escherichia coli - conformation 3. https://www.ebi.ac.uk/pdbe/entry/emdb/EMD-12655

Kolata P, Efremov RG (2021). RCSB Protein Data Bank ID 7NZ1. Respiratory complex I from Escherichia coli focused refinement of cytoplasmic arm. https://www.rcsb.org/structure/7NZ1

Kolata P, Efremov RG (2021). RCSB Protein Data Bank ID 7NYH. Respiratory complex I from Escherichia coli focused refinement of membrane arm. https://www.rcsb.org/structure/7NYH

Kolata P, Efremov RG (2021). RCSB Protein Data Bank ID 7NYR. Respiratory complex I from Escherichia coli conformation 1. https://www.rcsb.org/structure/7NYR

Kolata P, Efremov RG (2021). RCSB Protein Data Bank ID 7NYU. Respiratory complex I from Escherichia coli conformation 2. https://www.rcsb.org/structure/7NYU

Kolata P, Efremov RG (2021). RCSB Protein Data Bank ID 7NYV. Respiratory complex I from Escherichia coli conformation 3. https://www.rcsb.org/structure/7NYV

---

**Figure 1 - figure supplement 1.**
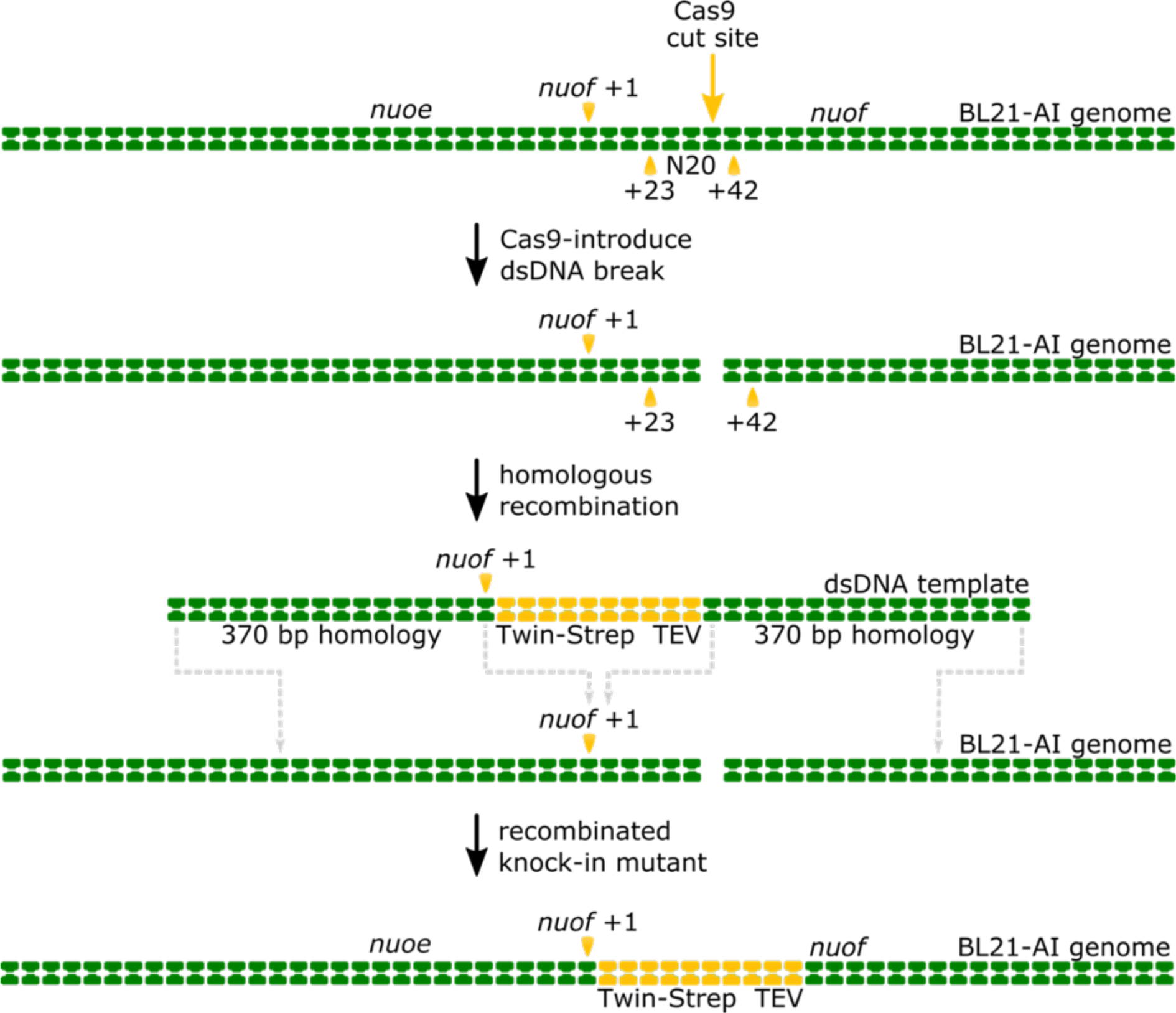
Schematic representation of Crispr-Cas9-enabled incorporation of the twin- strep tag into the N-terminus of the genomically-encoded NuoF subunit. The Cas9 enzyme introduces a double-stranded DNA (dsDNA) break into the nuof locus within the *E. coli* BL21-AI genome at the 20-nucleotide target sequence (N20), located 23 base-pairs (bp) downstream of the nuof +1 site. Subsequently, the λ-Red mediated homologous DNA recombination incorporates the supplied dsDNA template comprising the knock-in cassette flanked by 370 bp long homologous regions into the genome. The recombination results in insertion of sequences coding for the Twin-Strep tag and TEV protease recognition site right after the nuof transcription start site.

**Figure 1 - figure supplement 2.**
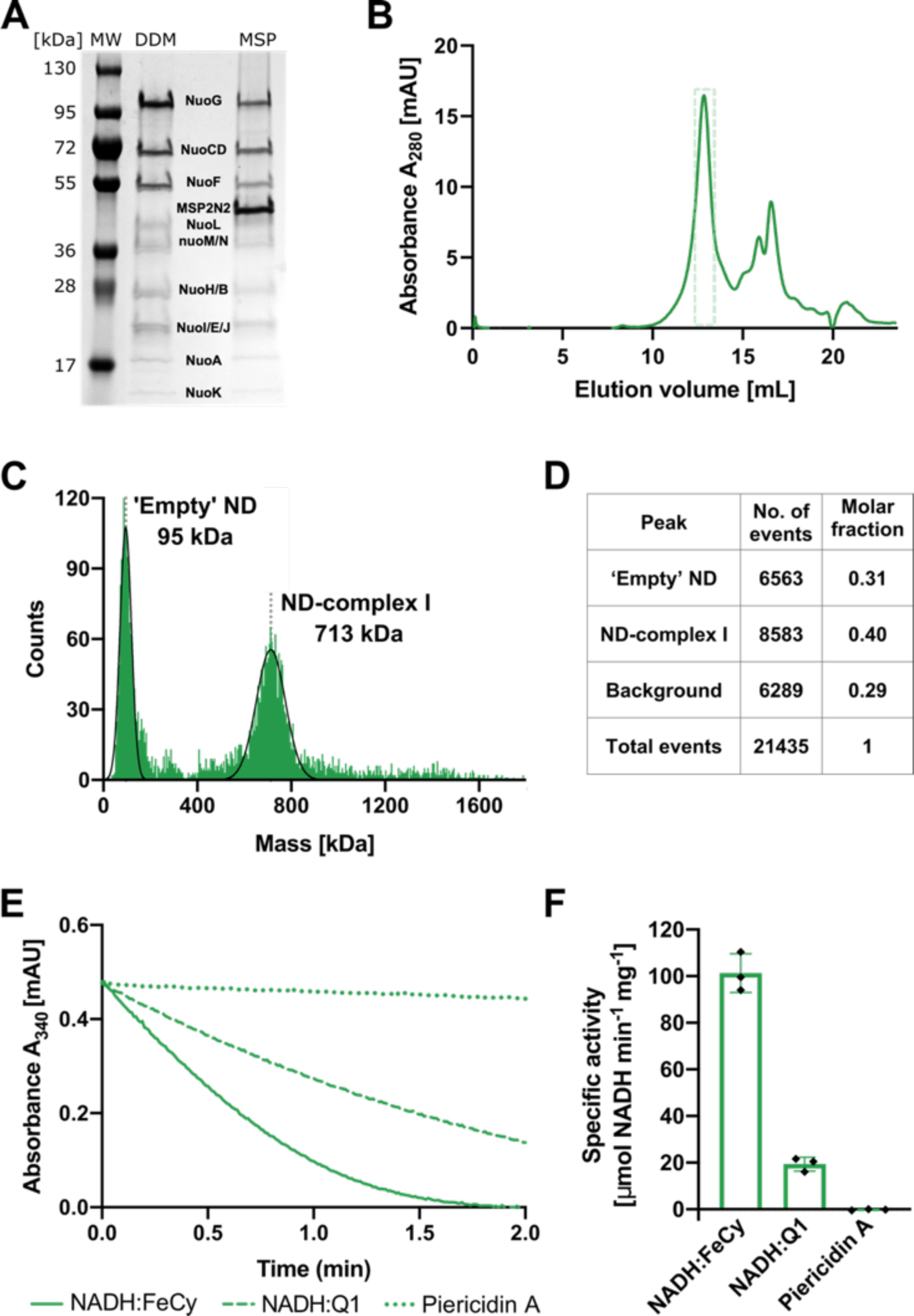
Purification and biochemical characterization of *E. coli* complex I reconstituted into lipid nanodiscs. (A) SDS-PAGE depicting the sample solubilized in detergent after affinity chromatography (DDM) and the sample reconstituted into lipid nanodiscs after size-exclusion chromatography (ND). (B) Size-exclusion chromatography profile after complex I reconstitution into lipid nanodiscs. The main peak at 13 mL consists mostly of the intact reconstituted complex I. Fractions marked by the green, dashed-rectangle were pooled, concentrated, and used for cryo-EM and activity assays. The second peak (retention volume 15 ml and higher) contains empty nanodiscs and the dissociated cytoplasmic arm. (C,D) Mass photometry of the reconstituted complex I pooled form the main gel filtration peak. (C) The representative mass histogram, showing two main peaks: ‘Empty’ nanodiscs at 95 kDa and the nanodisc-reconstituted complex I at 713 kDa. (D) Molar fractions of components identified in the histogram C. (E) Representative traces of the spectrophotometric activity assays: NADH:FeCy (solid line), NADH:Q1 (dashed line), and NADH:Q1 in the presence of 20 µM Piericidin A (dotted line). The concentration of complex I was 2.5 times lower for NADH:FeCy compared to that for the NADH:Q1 assay. (F) Values of Vmax for the three assay conditions described in panel (E). The graph shows mean ± SD, n=3. Individual measurement results are indicated as diamonds.

**Figure 1 - figure supplement 3.**
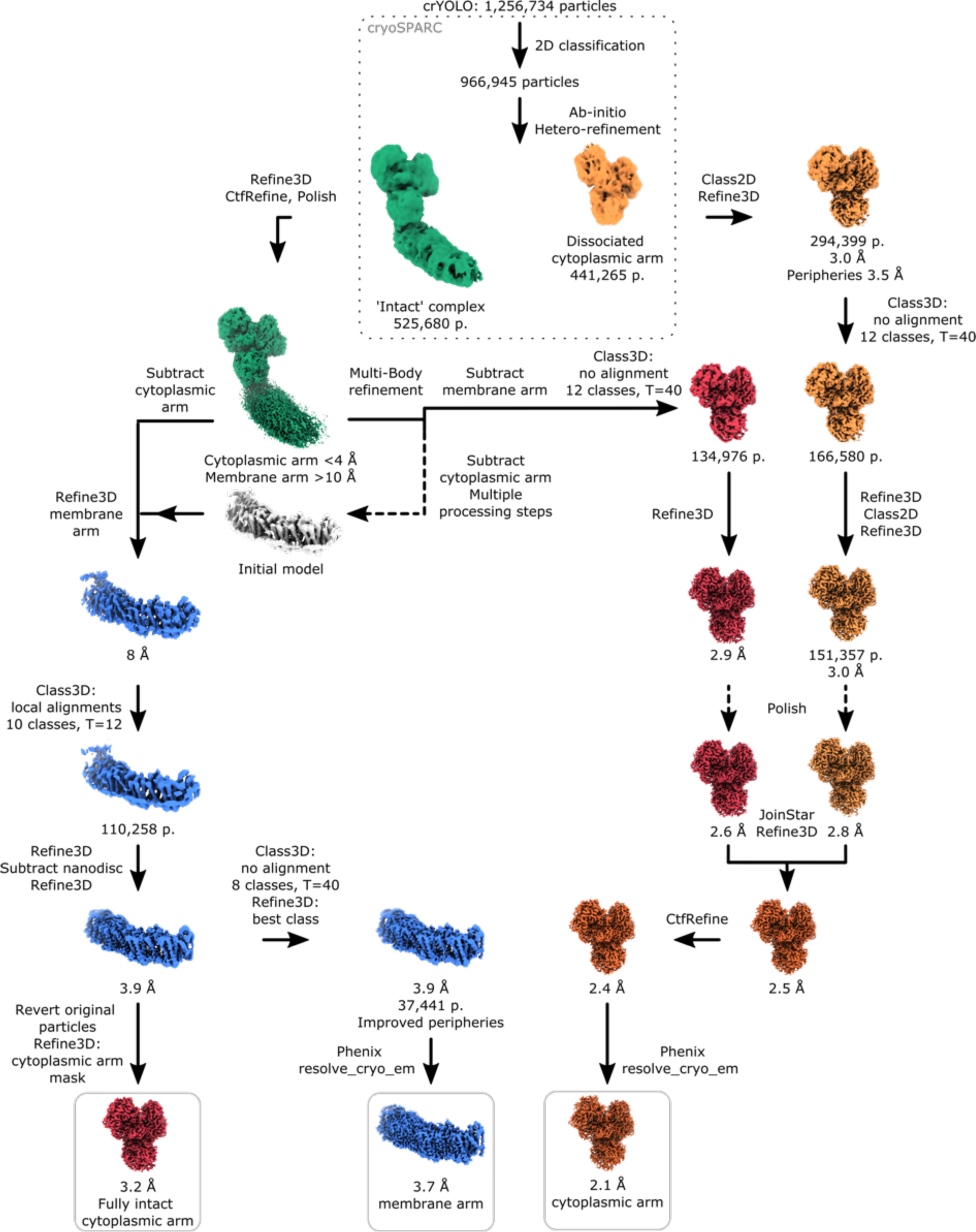
Image processing diagram. The scheme indicates the principal steps during image processing that resulted in reconstructions of the peripheral and membrane arms.

**Figure 1 - figure supplement 4.**
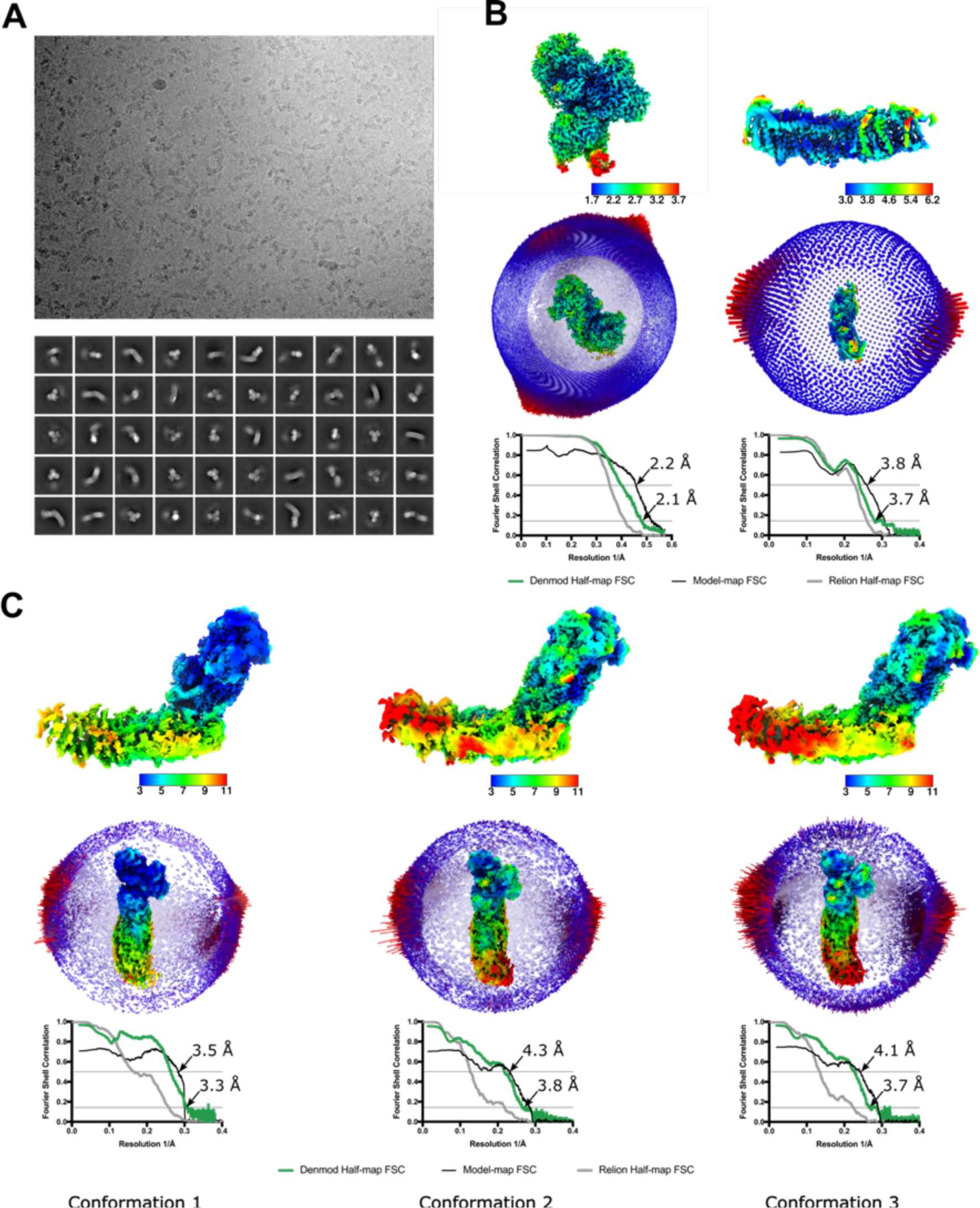
Properties of the cryo-EM sample and the final reconstructions. (A) A representative micrograph (top) and 2D classes of the entire complex I (bottom). (B,C) Local resolution maps, angular distribution, and FSC plots for reconstructions of the cytoplasmic (B, left) and membrane (B, right) arms as well as for the three conformations of the intact complex I (C).

**Figure 1 - figure supplement 5.**
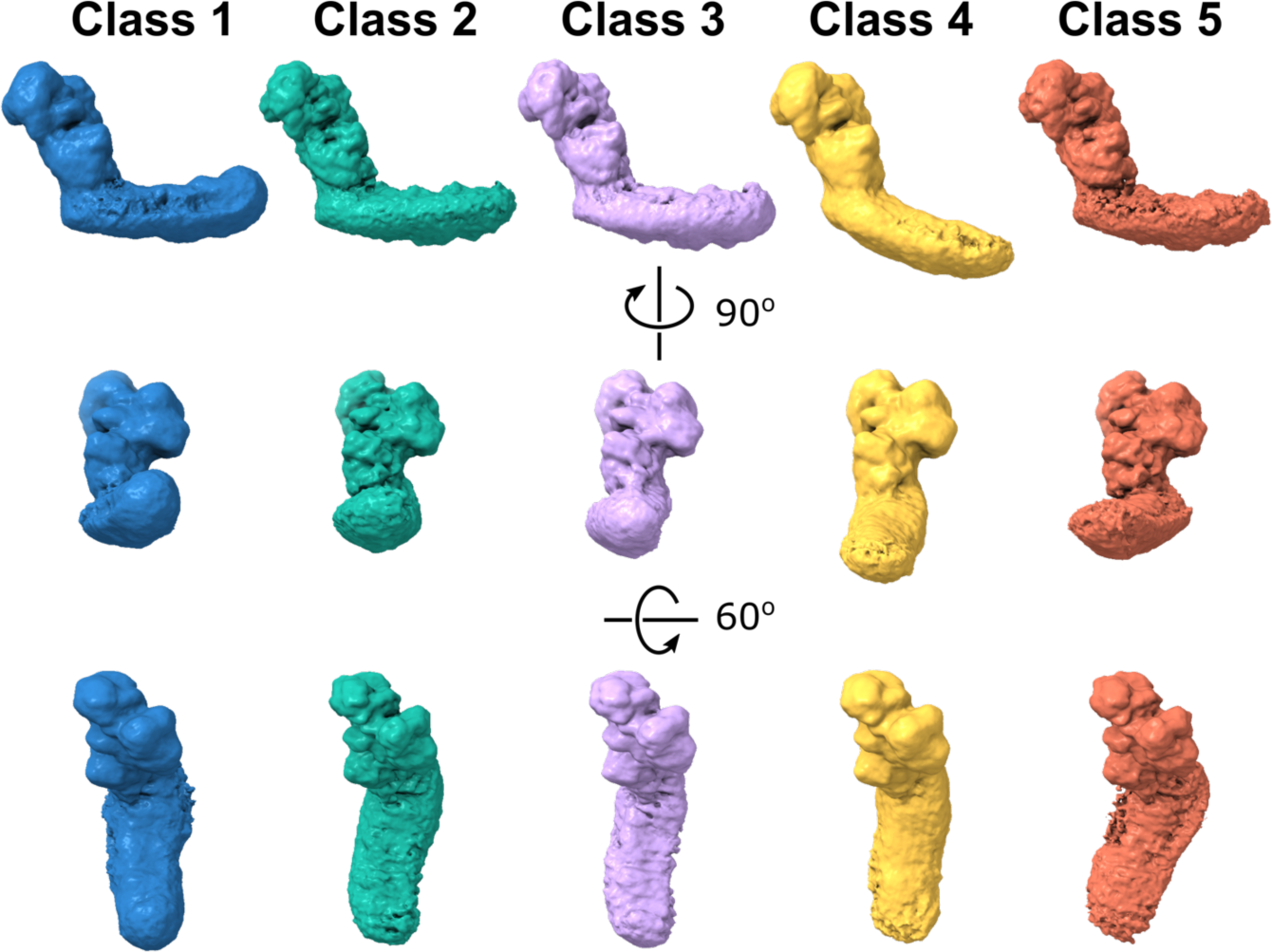
Dynamic connection between peripheral and membrane arms. Representative 3D classes show flexibility between the arms in the *E. coli* complex. The conformations were obtained by 3D classification of all intact complex I particles aligned to the peripheral arm.

**Figure 1 - figure supplement 6.**
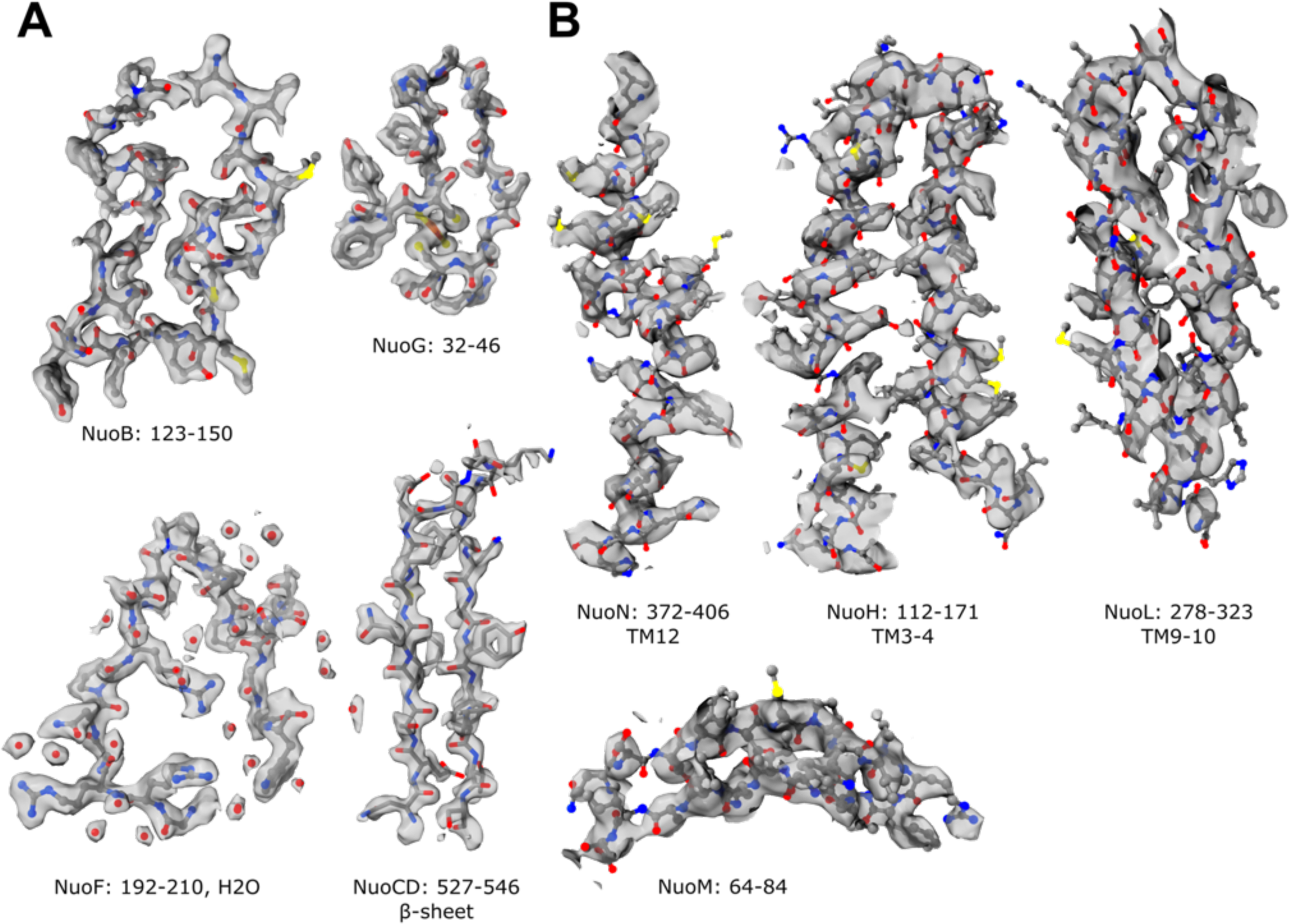
Representative cryo-EM map densities. Examples of density maps for the reconstructions of the (A) cytoplasmic and (B) membrane arm.

**Figure 2 - figure supplement 1.**
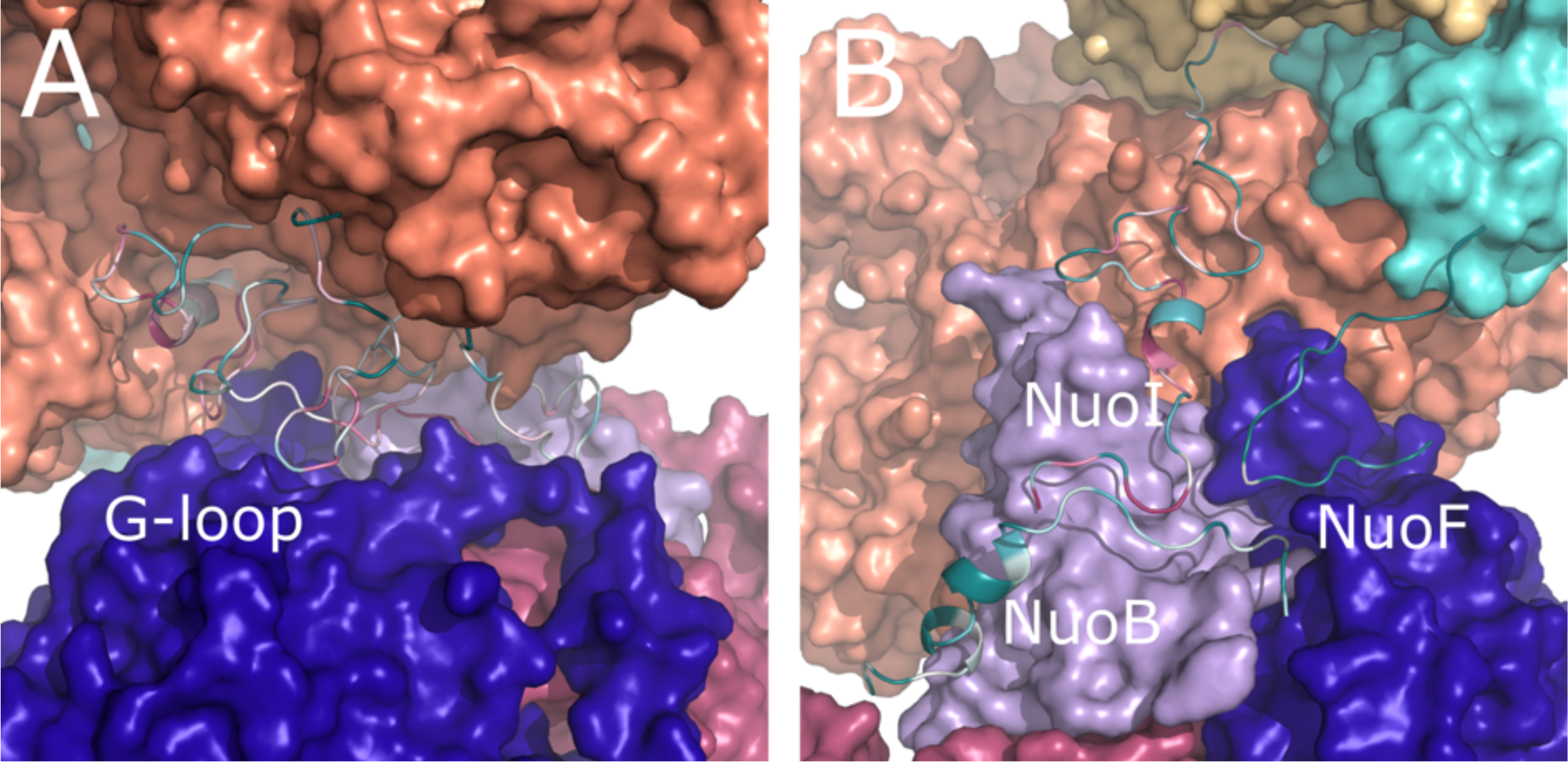
Conservation of *E. coli*-specific tails in the peripheral arm subunits. (A) Insertion into subunit NuoG, the G-loop. (B) C-terminal tails in subunits NuoF, NuoI and NuoB. Color coding of the subunits is the same as in Figure 1. Conservation was calculated using ConSurf server and is color coded from green to white to purple as the degree of conservation increases.

**Figure 3 - figure supplement 1.**
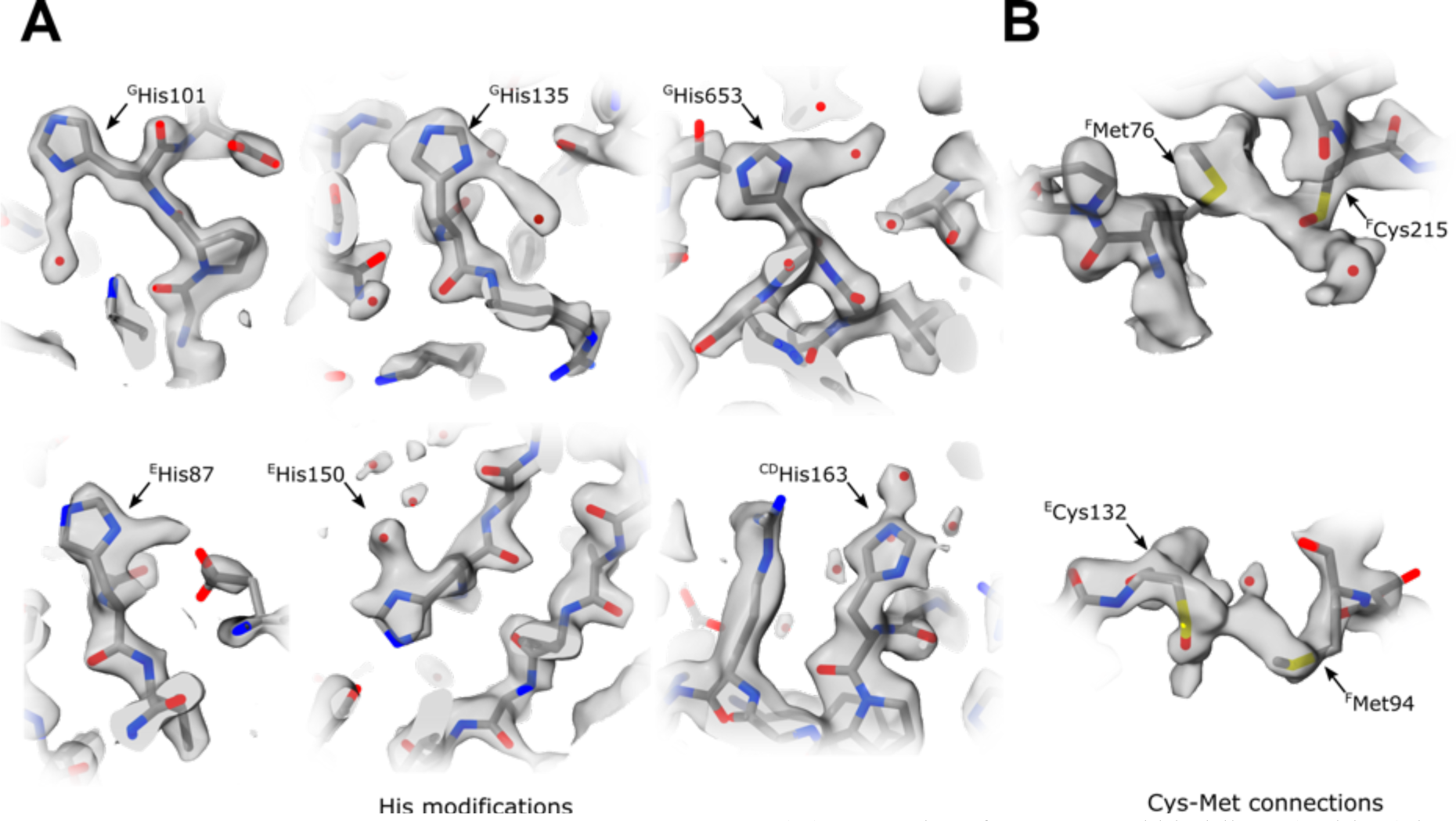
Unusual density features. (A) Several surface-exposed histidines (Table 6) have extended density protruding from the imidazole ring in the plane of the ring. These extensions may represent a tightly bound heavy atom, but these features are heterogeneous (length between 2 and 4 Å) and have a very heterogeneous chemical environment; therefore, they cannot be attributed to a single type of bound atom or modification. We tentatively modelled them as water molecules. (B) In three locations, density bridging sulfur atoms of surface-exposed and closely positioned cysteine-methionine couples are bridged by density with reproducible elongated features (Cys-Met: ^F^Met76-^F^Cys215, ^F^Met94-^E^Cys132, ^B^Met106-^B^Cys102). We could not assign the density to any known modification or tightly bound chemical present in the protein purification buffer. Because complex I was purified without reducing agents, the corresponding Cys residues were tentatively modelled as cysteines oxidized to sulfenic acid.

**Figure 4 - figure supplement 1.**
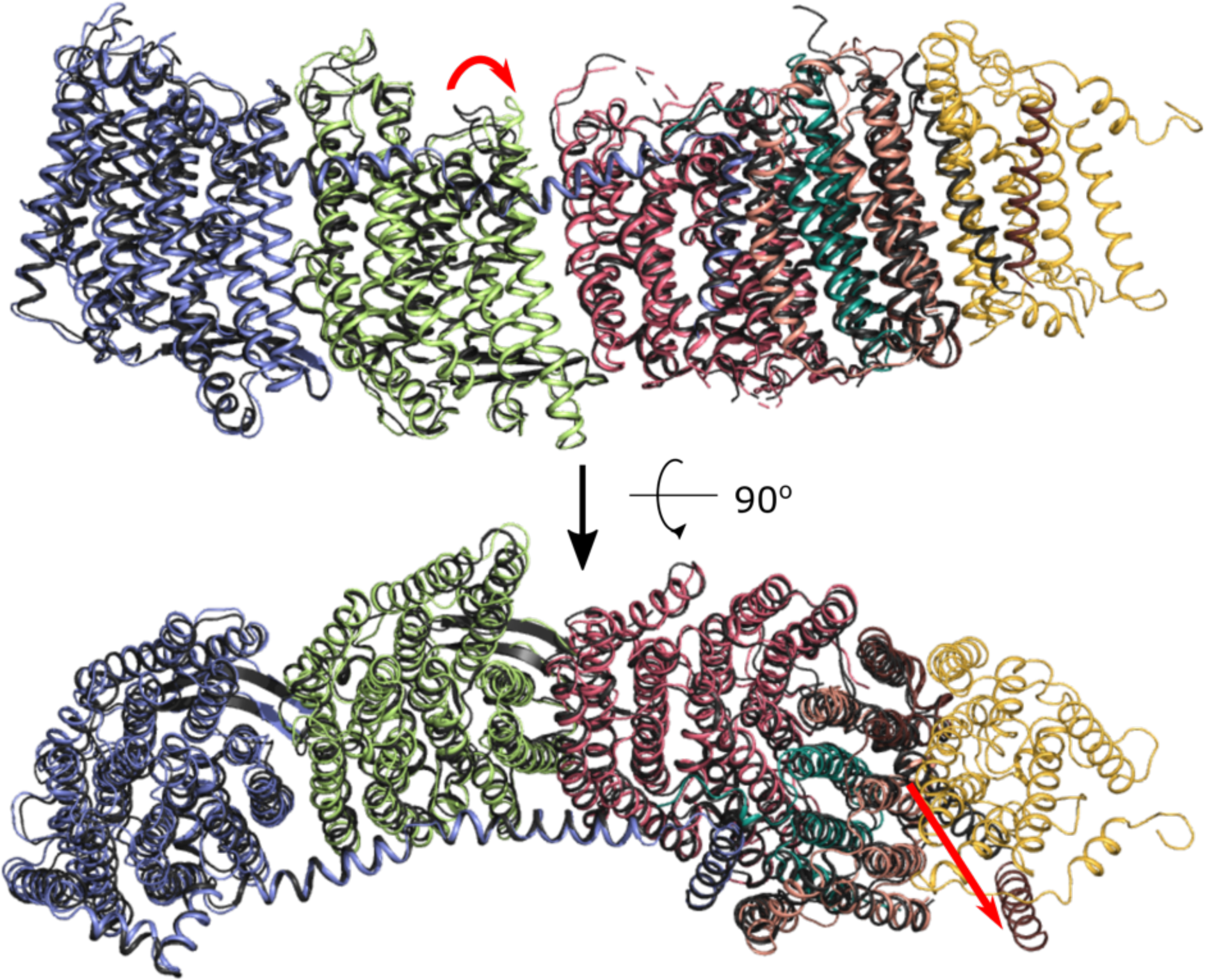
Comparison of X-ray and cryo-EM structures of the membrane domain. The cryo-EM structure is colored as in Figure 1, the X-ray model is shown in black. Significant shifts in the cytoplasmic loop of NuoM and the shift of ^A^TM1 are indicated with red arrows. Part of the b-hairpin observed in the crystal structure forms an extension of ^M^TM6a in the cryo-EM structure. This happens despite this loop not being involved in direct crystal contact.

**Figure 4 - figure supplement 2.**
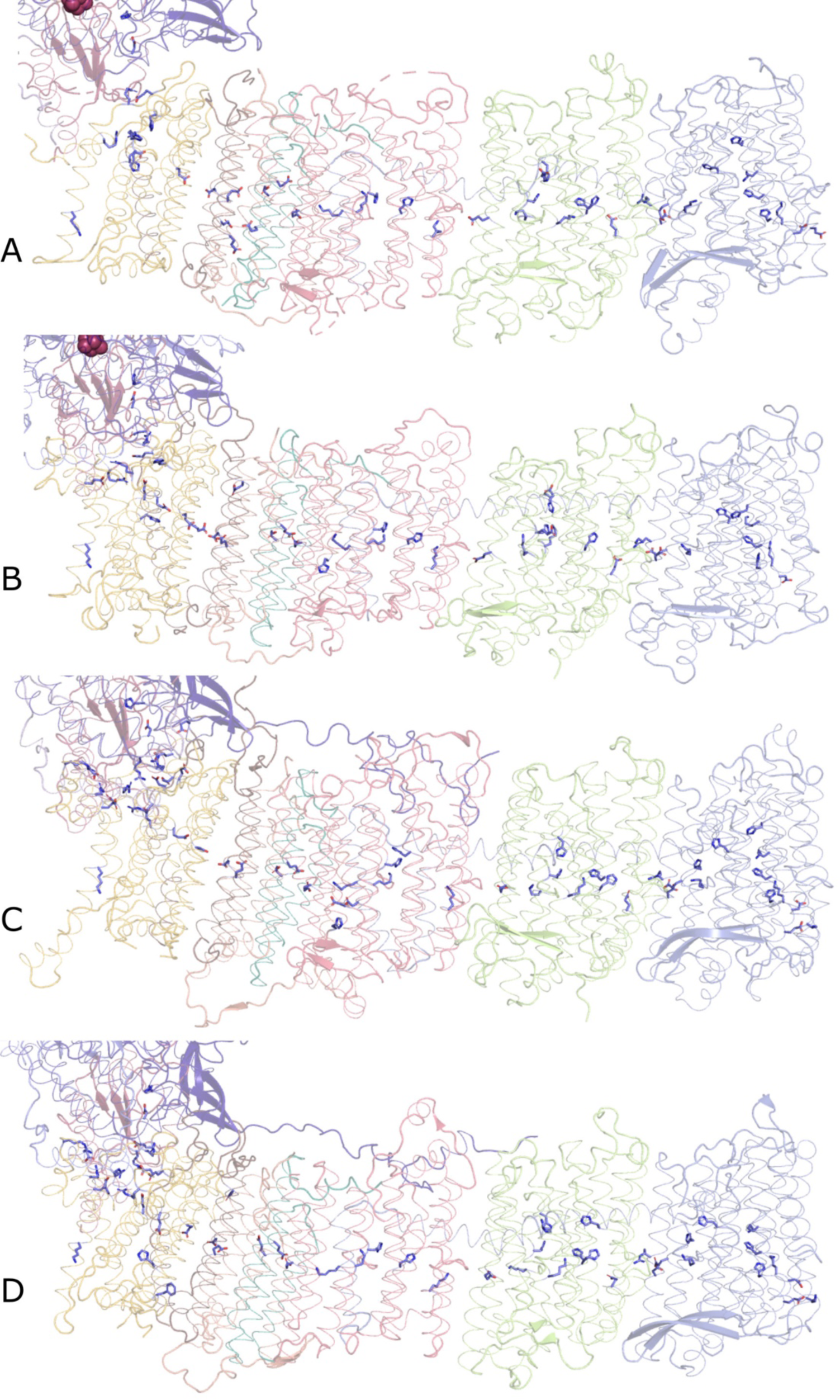
Conserved chain of ionizable residues. Comparison of the chains of ionizable residues lining the Q-cavity and positioned in the hydrophobic region of the lipid membrane in (A) *E. coli*, (B) *T. thermophilus*, (C) *Yarrowia lipolytica,* and (D) *Ovis aries* complex I. In C and D, only the core subunits are shown.

**Figure 5 - figure supplement 1.**
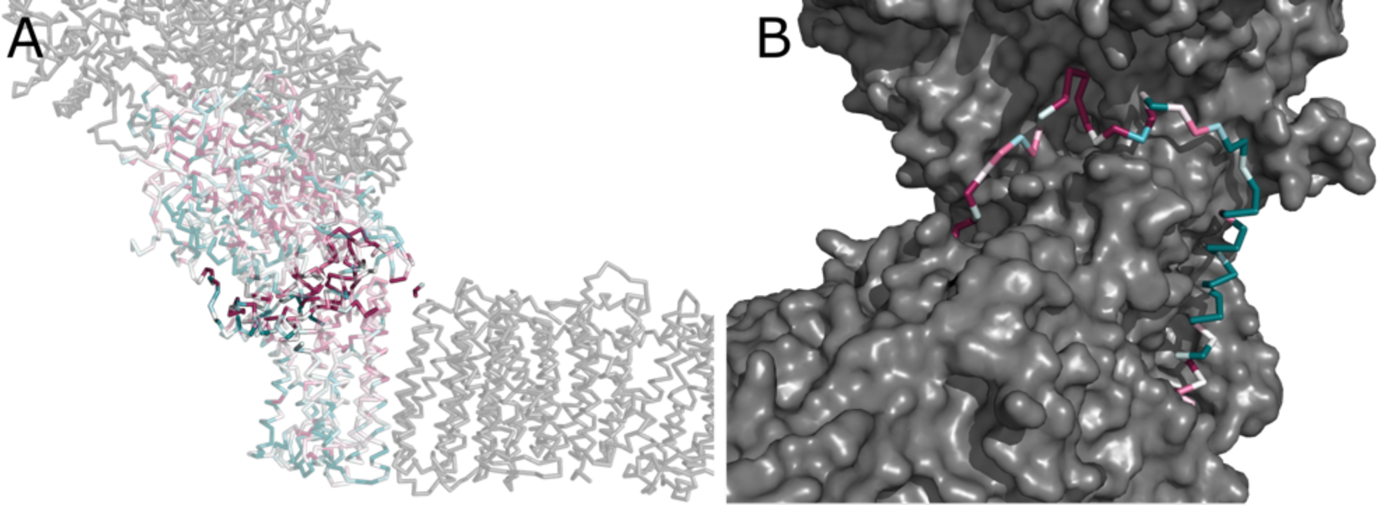
Conserved interface between the arms. (**A**) Conservation of subunits contributing to the interface between the peripheral and membrane arms, calculated in ConSurf and color-coded from green to white to purple as the degree of conservation increases. Residues directly contributing to the interaction between arms are highlighted. (B) The conserved region of the NuoA TMH1-TMH2 loop forms a plug, which fills a crevice between the subunits NuoD, NuoB, and NuoH.

**Figure 5 - figure supplement 2.**
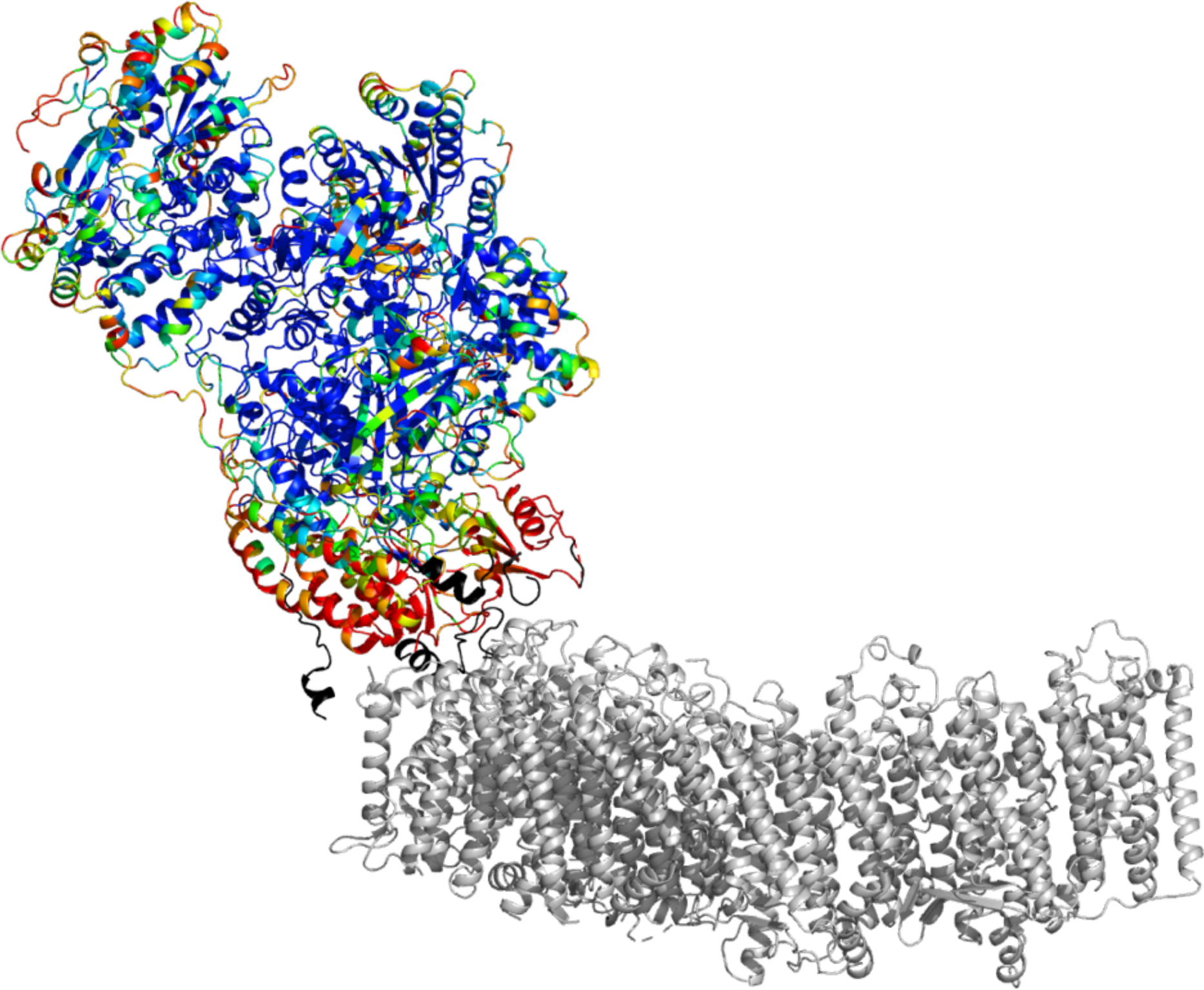
B-factors of the peripheral arm show higher mobility at the arms interface. The cartoon representation is shown. The peripheral arm is colored by B-factors. The rainbow color palette scales from 5 Å^2^ - blue to 100 Å^2^- red. Regions completely disordered in the map of focused reconstruction of the peripheral arm but showing density in in the reconstructions of the intact complex are colored in black. The membrane arm is shown in grey for reference.

Movie 1 Composite density map of *E. coli* complex I is shown along with density of the lipid nanodisc. The homology model of TMH1is shown in ribbon representation.

Movie 2 The fragmented density of the NuoM TM8 is surrounded by well-resolved TMHs.

